# Using visual scores and categorical data for genomic prediction of complex traits in breeding programs

**DOI:** 10.1101/2023.02.27.530308

**Authors:** Camila Ferreira Azevedo, Luis Felipe Ventorim Ferrão, Juliana Benevenuto, Marcos Deon Vilela de Resende, Moyses Nascimento, Ana Carolina Campana Nascimento, Patricio Munoz

## Abstract

Most genomic prediction methods are based on assumptions of normality due to their simplicity, robustness, and ease of implementation. However, in plant and animal breeding, target traits are often collected as categorical data, thus violating the normality assumption, which could affect the prediction of breeding values and the estimation of crucial genetic parameters. In this study, we examined the main challenges of categorical phenotypes in genomic prediction and genetic parameter estimation using mixed models, Bayesian approaches, and machine learning techniques. We evaluated these approaches using simulated and real breeding data sets. Our contribution in this study is a five-fold demonstration: (i) collecting data using an intermediate number of categories (1 to 3 and 1 to 5 scores) is the best strategy, even considering errors and subjectivity associated with visual scores; (ii) in the context of genomic prediction, Linear Mixed Models and Bayesian Linear Regression Models are robust to the normality violation, but marginal gains can be achieved when using Bayesian Ordinal Regression Models (BORM) and Random Forest Classification technique; (iii) genetic parameters are better estimated using BORM; (iv) our conclusions using simulated data are also applicable to real data in autotetraploid blueberry, which can guide breeders’ decisions; and (v) a comparison of continuous and categorical phenotype testing for complex traits with low heritability, found that investing in the evaluation of 600-1000 categorical data points with low error, when it is not feasible to collect continuous phenotypes, is a strategy for improving predictive abilities. Our findings suggest the best approaches for effectively using categorical traits to explore genetic information in breeding programs, and highlight the importance of investing in the training of evaluator teams and in high-quality phenotyping.

**Key message:** An approach for handling categorical data with potential errors and subjectivity in scores was evaluated in simulated and blueberry recurrent selection breeding schemes to assist breeders in their decision-making.

## Introduction

Over the last century, plant and animal breeders have used quantitative genetics to estimate genetic parameters and predict phenotypic traits. These traits are typically modeled as a function of the genetic makeup of plants (genotype) and the conditions in which that plant developed (environment). This traditional framework relies on one critical assumption: the normality of phenotypic traits. The use of linear models for phenotypes that follow a Normal (or Gaussian) distribution is attractive due to its simplicity, robustness, straightforward implementation, and support by a well-established theory. However, data sets in plant breeding often fall outside the scope of normality, which offers additional statistical challenges. In these cases, a central question faced by biometricians is how they should model non-Gaussian phenotypes, and if such normality violation may affect estimates of genetic parameters of key interest.

A core assumption underlying the use of linear models is the normality of the residuals and, consequently, the response variables. Data sets in animal and plant breeding are often collected in the form of binary (e.g., the infection status of diseases), proportions (e.g., mortality rates), or counts (number of emerging seedlings or number of fruits) forms. At least, in theory, violation of the normality assumption in these data sets can invalidate the model and affect future decisions, such as leading to highly imprecise estimates (Schielzeth et al., 2020).

To address these issues, breeders and biometricians have used various strategies. The simplest approach is ignoring the lack of normality, under the argument that large sample sizes follow the central limit theorem – which states that treatment means have an approximately normal distribution when sample sizes are large enough (Montesinos-López, Montesinos-López, and Crossa, 2017). Another common alternative is transforming the phenotypes, which can stabilize the residual variation, and hence help fulfill the assumptions required by linear modeling. Although both alternatives are popular in the plant breeding literature, there is empirical evidence showing that the accuracy and power of statistical inference can be reduced when shoehorning the data into classical statistical methods (Stroup, 2015).

A more formal approach to non-Gaussian traits relies on the use of a generalized linear model (GLM) (McCullagh and Nelder, 1989). In this model, the mean of a response is modeled as a function of explanatory variables, and the response variable is assumed to be conditionally distributed, according to an exponential family distribution (e.g., Binomial, Poisson, or Gamma distributions for trial, count, or strictly positive real responses, respectively).

Another important aspect is that the breeders’ model phenotypic observations are a function of variations at the DNA level. Referred to as genomic selection, this methodology is a form of marker-assisted selection in which all available molecular markers are used to predict quantitative traits (Meuwissen, Hayes, and Goddard, 2001). Despite its importance, the debate around using of non-Gaussian traits remains the same: most genomic prediction models are based on linear regression models that assume continuous and normally distributed phenotypes, without clear evidence on the impact of normality violation on estimating genetics parameters.

Some recent studies have relaxed these assumptions and applied threshold models and Bayesian ordinal regression. For example,Montesinos-López, Montesinos-López, Pérez-Rodríguez, Eskridge, et al. (2015) introduced genomic selection models for ordinal traits in maize, and reported gains when genotype-by-interaction was taken into account. They also reported the use of ordinal logistic regression for predictions, under the argument that ordinal models are more robust for dealing with outlying data and provide interpretable results (Montesinos-López, Montesinos-López, Pérez-Rodríguez, Campos, et al., 2015). In animal science, ordinal and continuous data were compared, and the use of threshold traits resulted in markedly lower accuracy than a linear model (Kizilkaya, Fernando, and Garrick, 2014). More recently, Merrick et al. (2022) reported that using machine learning methods led to higher predictive accuracy for the classification and prediction of traits with skewed distributions.

In this context, it is still unclear how to analyze phenotypic data that are naturally normally distributed, but are categorized to simplify assessments and reduce the costs of phenotyping. A prime example is yield evaluation in fruit crops. Using blueberry as our biological model, yield is commonly evaluated after all berries are manually picked and weighed. Remarkably, harvesting takes place multiple times during the crop season, which makes the process slow, laborious, and expensive. As an alternative, breeders visually classify the genotypes using visual scores that can range between 1(low production) to 5 (high production). Similarly, plant pathologists also have relied on visual scores to screen quantitative diseases, but use scores ranging from 1 (low infection) to 9 (high infection). Even with the recent progress of computer vision and phenomics, it is still labor-intensive to carry out such quantitative assessments in the laboratory, when compared to field conditions. However, the use of visual assessments does not relieve breeders of important decisions, i.e. the choice of increased costs of screening a large population using visual scores or reduced experimental accuracy using numerical scoring for continuous variation in a variable.

Aiming to shed light on the relevance of using visual and categorical traits in plant breeding, we conceptualized this study in two sections. First, we simulated continuous data, with Gaussian distribution, categorized it, and included different levels of noise – mimicking potential errors associated with recording visual scores. We examined the impact of using categorical traits for prediction and genetic inference using parametric and non-parametric models. Secondly, we used data collected in a real blueberry breeding population (our biological model) and modelled categorical traits evaluated over multiple years and locations by the University of Florida (UF) breeding program. Collectively, in this study we addressed the following questions: (i) what is the best strategy for recording and modelling categorical data? (ii) what is the effect of the operators’ experience (error level) when estimating genetic parameters? (iii) should non-Gaussian traits be modelled using parametric or non-parametric methods? and finally, (iv) how can breeders allocate resources for phenotyping to collect continuous and categorical data, to maximize predictive gains?

## Methods and Materials

The Materials and Methods section of the study is organized as follows. The “Simulated Data” section describes the development of simulations for the experiments, using a stochastic model that considers various levels of noise/errors. The “Data analysis” and “Scenarios of Analyses” sections outline the use of parametric and non-parametric models for predicting and estimating genetic parameters. At this stage, no real data is included. The “Genomic prediction and efficient measures calculation” section presents the metrics used to compare the different analysis approaches. Finally, the “Real Data Analyses” section applies the findings from the simulated results to real data analyses on blueberries.

### Simulated Data

The effect of categorizing continuous traits using a simulated data set was first studied by simulating phenotypic traits with contrasting genetic architectures. The simulated genome consisted of 10 pairs of chromosomes with a genetic length of 1.43 Morgans. Sequences for each chromosome were randomly chosen to have 1,000 causal loci per chromosome (a total of 10,000 across the genome) and were generated using the Markovian Coalescent Simulator (Chen, Marjoram, and Wall, 2009). After generating genome sequences, we created 50 founder genotypes that were used as initial parents in the burn-in phase. The following steps involved crosses between highly heterozygous hybrids. After simulating the crosses, 10% of the resulting F1 progenies (5,000 genotypes) were selected based on their estimated breeding value (500 genotypes). Then, using genomic models, we selected 50 genotypes to be parents in the next breeding cycle. This simulation represents a typical small effective population size (Ne = 50). The methods and analysis scenarios were evaluated using the average of ten replicates, and each replicate consisted of: i) a burn-in phase that consisted of 20 cycles of breeding, and ii) an evaluation phase that simulated future breeding with different analysis strategies. In this way, we simulated a classical recurrent selection breeding program in which the allele frequencies of target traits are increased by selecting the best individuals and crossing them to create a new generation.

In our breeding design, we simulated traits with two genetic architectures: (i) a qualitative trait controlled by three large quantitative trait loci (QTL) and high heritability (0.60); and (ii) a quantitative trait following the infinitesimal model, with 100 QTL and low heritability (0.10). For each QTL, we assigned one additive effect on the phenotype following a normal distribution with zero mean and variance, resulting in the desired heritability level (Gaynor, Gorjanc, and Hickey, 2021). We added a random deviation from a normal distribution N(0,100) to the genotypic value. All simulations were carried out using AlphaSimR (Gaynor, Gorjanc, and Hickey, 2021), following a similar crossing and selection design as those described by Batista et al. (2021).

As part of the simulation process, continuous phenotypic traits (*Y*_*i*_) were categorized as follows: two (1 to 2), three (1 to 3), five (1 to 5), and nine (1 to 9) categories. The use of four different numbers of categories mimics visual scores which are commonly used in plant breeding programs. To create these categories 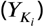, we used the following thresholds to categorize the simulated phenotype values:

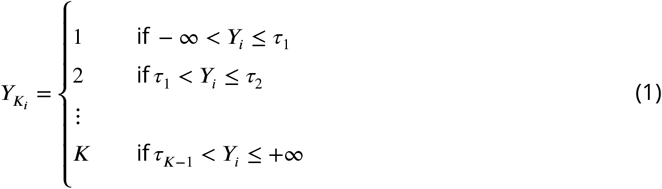

where *τ*_1_, *τ*_2_, …, *τ*_*K*−1_ are the non-equidistant thresholds based on quantiles distribution and K is the number of categories (*K* = 2, 3, 5 or 9).

Assuming that visual scores may be subject to errors due to the experience of the person recording them, we also simulated different levels of noise, or error. We considered three levels of error: low, moderate, and high, which caused 20%, 50%, and 70% misclassification in the scores, respectively. This approach is similar to what has been reported in genomic studies that have evaluated the effect of mislabeling on genomic prediction trials (Biffani et al., 2017; Yabe, Iwata, and Jannink, 2018). Importantly, an error rate of 20% –even considered as “low” in this study – is still a very conservative value for most breeding programs. Figure 1 summarizes the different categorical classes and noise levels.

**Figure 1.**
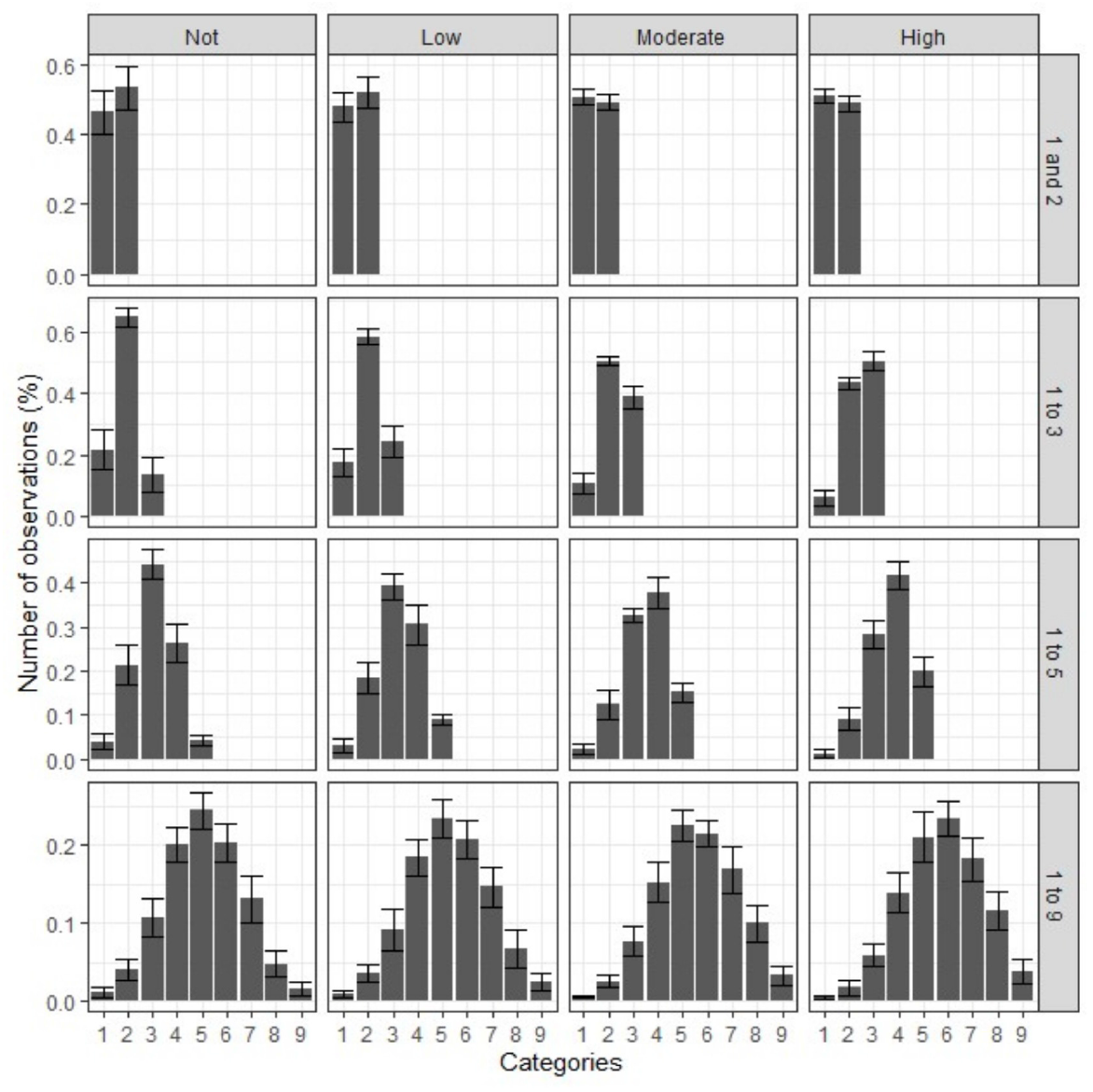
Distribution of observations simulated by four different numbers of categories, considering three noise levels (low – 20% of misclassification, moderate – 50% of misclassification, and high – 70% of misclassification) and no errors.

### Data analyses

For data modeling, we employed two main approaches: statistical methods (including Generalized Linear Mixed Model, Linear Mixed Model, Bayesian Ordinal Regression Model, and Bayesian Linear Regression Model) and machine learning methods (Random Forest Regression and Random Forest Classification).

### Generalized linear mixed model

The Generalized Linear Mixed Model (GLMM) is defined as:

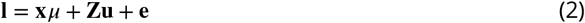

where **I** is the vector of latent variables (called liabilities) for the vector of phenotypic values represented by y, **x** is a vector of the same dimension of **l** being all elements equal to 1, *μ* is the trait mean, **u** is the vector of additive genetic random effects of individuals with 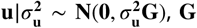 is the additive genomic relationship matrix (VanRaden, 2008) between individuals and 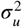 is the additive genetic variance, **Z** is the matrix of incidence of random effects and **e** is the vector of random errors. In a generalized linear model, it is assumed that each observation of the variable Y has a distribution belonging to the exponential family. In this case, the expected value of Y is defined as:

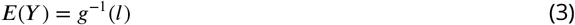

where *g*(.) is the link function. The linear mixed model (LMM) is a particular case of GLMM, in which the variable Y follows a normal distribution with mean **x***μ* +**Zu** and a covariance matrix given by **I***σ*^2^, where *σ*^2^ is the residual variance, and whose link function is the identity (McCullagh and Nelder, 1989). Suppose the Y variable consists of levels of some categorical factors and has a natural ordering, an adequate link function is the probit function. Ordinal categorical predictions for phenotypes with K categories are defined based on threshold parameters *γ*′ = (*γ*_**0**_ = −∞, *γ*_**1**_, *γ*_2_, …, *γ*_**K**−**1**_, *γ*_**K**_ = +∞) that have a continuous scale and relate to the observed ordinal categorical response, according to:

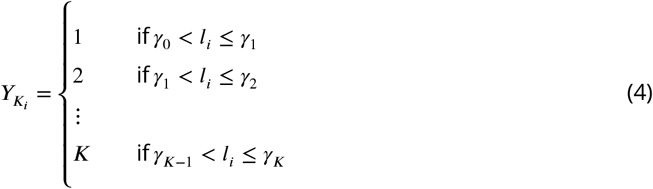

where *i* = 1, …, *n* and *n* is the number of the observations.

Therefore, in the ordinal model, *Y*_*i*_ is a random variable that takes values 1, …, *K*, with the following probabilities:

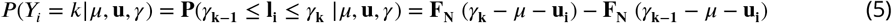

where *F*_*N*_ is the Gaussian cumulative distribution function.

A procedure for Henderson’s generalized linear mixed model equations (GLMME) that leads to the BLUP in linear mixed models (Resende et al., 2018) can be defined as:

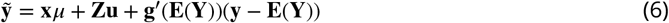

where 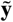 is observational variable adjusted and *g*′(.) is the first derivative of *g*(.). So, the estimation and prediction algorithms for the linear mixed model case can be adapted as the mixed model equations below:

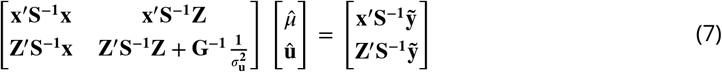

where **S**^−**1**^ is a diagonal matrix with elements equal to 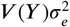 and 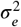 is the residual variance on the continuous scale (liabilities). The Restricted maximum likelihood (REML) estimators are given by:

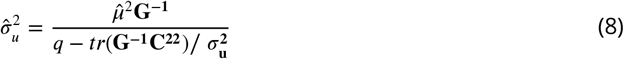

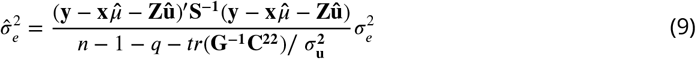

where *n* is the number of the observations, q is the number of random effect levels, tr is trace matrix operator, 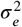 and 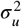 are values obtained in the previous iteration of the algorithm and *C*^22^ is the partition of the inverse of the coefficients of the mixed model equations, referring to random effects.

And the solution to the GLMME may be expressed in the following way:

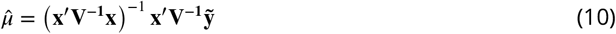

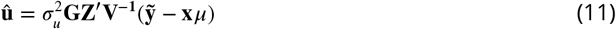

where 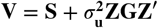.

These procedures were implemented in the ASReml package in software R (Butler, 2022), for each replicated data set within the scenarios and real data set.

### Bayesian Ordinal Regression Model

Model 2 can be used under a Bayesian framework. The Bayesian Ordinal Regression Model (BORM) model assumes the following prior distribution for the unknown parameter vector 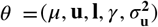:

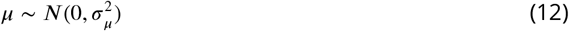

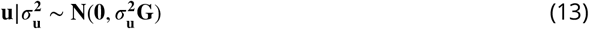

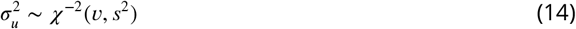

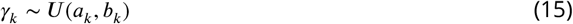

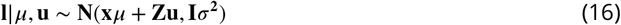

where **I** is an identity matrix and *σ*^2^ = 1 to reach identifiability for unobservable liabilities (even when the number of unknown parameters is higher than the sample size), 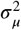 is a value assumed as 10^10^ that represents a vague prior knowledge, **G** is the additive genomic relationship matrix (VanRaden, 2008) between individuals and 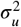 is the additive genetic variance. The *ν, s*^2^, *a*_*k*_ and *b*_*k*_ are called hyperparameters.

The joint posterior density of *μ*,**u**,*γ*,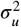 and the liabilities **l** is given by:

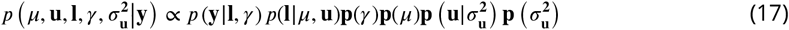

The inference of the parameters (*μ*, **u, l**, *γ*, 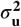) is based on their marginal posterior distributions, obtained indirectly from full conditional posterior distributions through the Markov Chain Monte Carlo (MCMC) algorithms. These full conditional posterior distributions were presented by López, López, and Crossa (2022). These procedures were implemented by the BGLR package in software R (Pérez and Campos, 2014) to each replicated data set within each scenario and to the real data by defining 500,000 iterations for the MCMC algorithms, a burn-in period of 50,000 MCMC cycles, and thin equals to 10 before saving samples from each, totaling 45,000 MCMC cycles. The **û**, 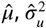 and 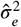 estimates were obtained as the posterior mean of their respective marginal posterior distributions. In the Bayesian Linear Regression Model (BLRM), the joint posterior density is simplified to

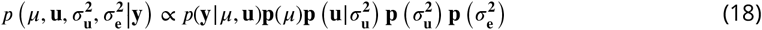

because the own response variable Y assumes the Normal distribution, e.g., 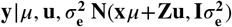.

### Random Forest Regression and Random Forest Classification

Random Forest Regression (RFR) and Random Forest Classification (RFC) are supervised machine learning methods based on tree algorithms that can apply to continuous, binary and categorical variables, respectively (James et al., 2013). The tree algorithms divide the predictor space (*M*_1_, *M*_2_, …, *M*_*p*_ – in this study, molecular markers) into several non-overlapping regions (*R*_1_, …, *R*_*J*_), and these stratifications are based on the optimization of cost functions. The regression tree is indicated for continuous traits, and the goal is to find boxes *R*_1_, …, *R*_*J*_ that minimize the Residual Sum of Squares given by:

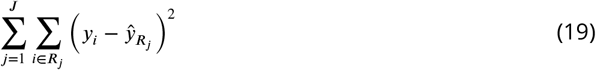

where 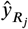 is the mean of response observations (continuous phenotypes) within the jth region. However, instead of considering each possible partition of space in J regions to reduce the computational time of the analyses, recursive binary splitting is performed by selecting the predictor *M*_*j*_ and the cutpoint s and then minimizing the equation:

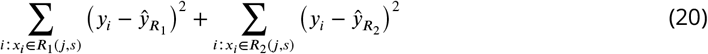

where 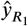 and 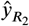 are, respectively, the mean of response observations in *R*_1_(*j, s*) and *R*_2_(*j, s*).

The classification tree procedure is very similar to a regression tree, but is indicated for binary and categorical traits, and uses other cost functions like the Gini index, given by:

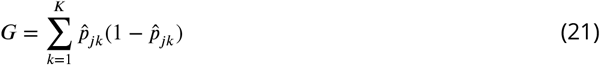

where 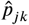 is the proportion of observations in the jth region that are from the kth category. Typically, a single regression or classification tree has high variability in its predictions and, as a result, a reduced ability to make accurate predictions. To improve the predictive and classification performance of the tree, refinements such as Random Forest can be used. Random Forest builds multiple trees that are decorrelated by using a subset of predictor variables in each partition, and then averages or takes the mode of the predicted values. This results in independent predicted values, which reduces the variability of the tree (Ho, 1995). The number of predictors (m) suggested by Hastie, Tibshirani, and Friedman (2009) is 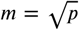 for classification trees (RFC) and *m* = *p*/3 for regression trees (RFR).

These procedures were implemented by the earth package in software R (Milborrow, 2021) to each replicated data set within each scenario and real data.

### Scenarios of analysis

For statistical purposes, genomic prediction was performed according to three theoretical scenarios:

- Continuous (CONT - benchmark): The simulated phenotypes (*Y*_*i*_) were used as the response variable in the Linear Mixed Model, Bayesian Linear Regression Model, or Random Forest Regression. This represents the benchmark scenario, where the true continuous distribution of the phenotypes is considered and analyzed with an appropriate model and probability distribution.
- Categorical-Continuous (CAT-CONT - practical): After categorizing the continuous phenotypes 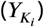, we decided to process the data into classical statistical methods, and used the ordinal response variable in the Linear Mixed Model, Bayesian Linear Regression Model or Random Forest Regression. This scenario represents what is typically done in breeding programs, where the phenotypes are collected as categorical but analyzed as continuous.
- Categorical (CAT - formal): After categorizing the continuous phenotypes 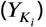, we used the categorical response variable into a Generalized Linear Mixed Model, Bayesian Ordinal Regression Model or Random Forest Classification. This scenario represents a formal statistical procedure for analyses of categorical phenotypes.

For a graphic representation of the “Simulated Data”, “Data Analyses” and “Scenarios of Analyses” sections, see Figure 2.

**Figure 2.**
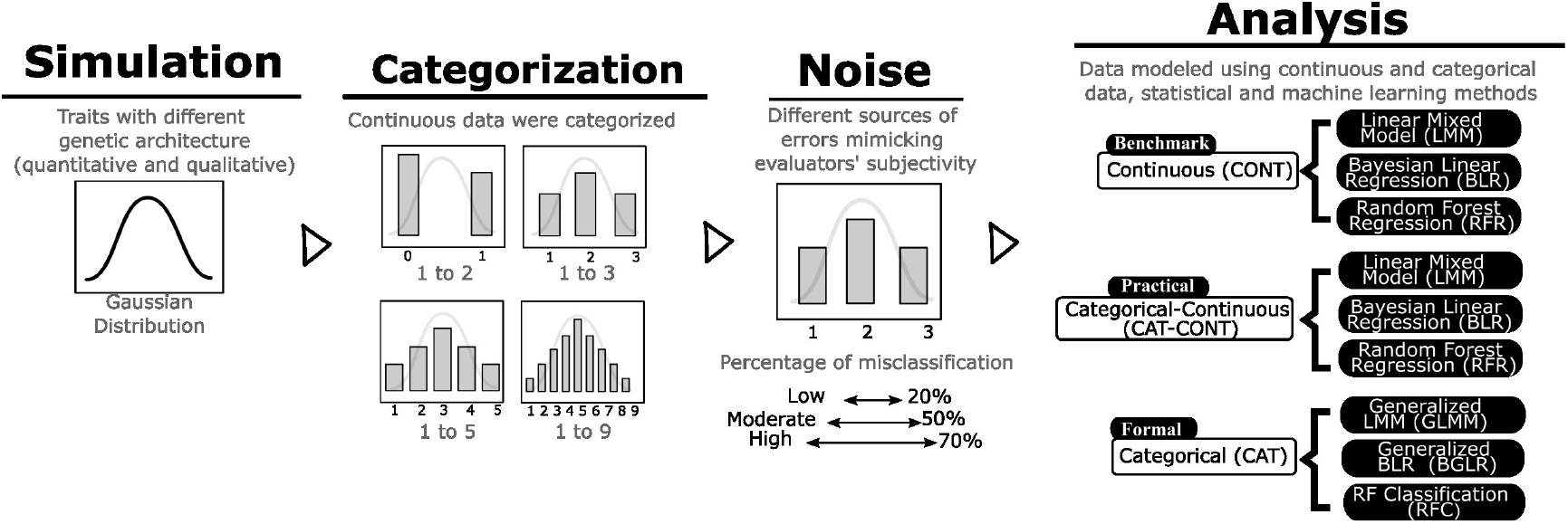
The following flowchart illustrates the procedures used in this study. The simulation component involves simulating traits with two genetic architectures (quantitative and qualitative) and Gaussian distributions. The categorization component involves delineating the continuous phenotypic traits into categories (1 to 2, 1 to 3, 1 to 5, and 1 to 9). The noise component includes the creation of noise levels (low – 20% of misclassification, moderate – 50% of misclassification, and high – 70% of misclassification) in the categorization, for mimicking the evaluator experience. The analysis component represents the application of parametric (Generalized and Linear Mixed Models, Bayesian Ordinal and Bayesian Linear Regression Models) and non-parametric (Random Forests Regression and Classification) methods to continuous and categorical traits. The CONT analysis scenario represents the benchmark scenario when continuous phenotypes are the response variables in the LMM, BLRM, and RFR. The CAT-CONT analysis scenario represents the typical practice in breeding programs where categorical phenotypes are the response variables in the LMM, BLRM, and RFR. The CAT analysis scenario represents a formal statistical procedure where the categorical phenotypes are the response variable in the GLMM, BORM, and RFC.

### Genomic prediction and efficient measures calculation

We compared the prediction performances using fivefold cross-validation and the following metrics: (i) accuracy, which is the correlation between the true breeding value (TBV) and genomic estimated breeding values (GEBV); (ii) predictive ability, which is the correlation between the phenotype and GEBV; (iii) prediction bias, which is the deviation from one of the coefficient of regression between TBV and GEBV; (iv) the percentage of agreement between the top 10% of individuals selected by each approach (CONT, CAT-CONT and CAT) and true breeding value; and finally (v) the correlation between the marker effects estimated by each approach.

We also computed genetic parameters in terms of heritability and selection gain. For the Bayesian approaches, heritability was computed for the kth value of the Markov chain of heritability, and is given by: 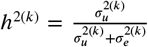, where 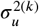 and 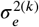 are values of the variance components of the kth iteration of the MCMC algorithm. Subsequently, the posterior mean was calculated. For the Henderson’s equations approach, the heritability is given by: 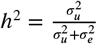, where 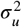 is estimated using REML, 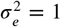 in the GLMM, and 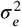 is estimated using REML in the LMM. For the random forest approach, the variances were calculated as being the variance of the GEBV 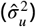 and residuals vector 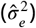 and then the heritability values were obtained by 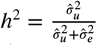.

Selection gain was calculated using the following expression, considering the selection of 10% of individuals by each approach: 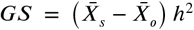, where 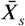 is the mean TBV of the selected population, 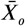 is the mean TBV of the original population, and *h*^2^ is the heritability estimate.

### Real Data Analyses

We extended our simulated analyses to real data, using Southern Highbush Blueberry genotypes from the University of Florida (UF). These genotypes consisted of advanced selections and interspecific hybrids. Briefly, as part of the recurrent selection strategy, the UF blueberry breeding program annually makes up to 200 crosses. These crosses are designed based on a combination of phenotypic, pedigree, and molecular information that predicts plant performance for yield, fruit quality, flavor, disease/pest resistance, and early season production. From crosses to a cultivar, a four-stage selection approach is used. In the first stage, 20,000 progenies are planted in high-density plantations from the approximately 200 crosses. The first evaluation cycle is conducted on one-year-old seedlings (Stage I), and about 10% of seedlings are advanced to the second stage (Stage II). In the second year, with more fruits available for evaluation, a new selection (10% of the approximately 2,000 remaining plants) is performed. Stage III consists of clonally propagated plants that have been established in a commercial field and evaluated in a 15-plant clonal plot. Of the genotypes in Stage III, approximately 10% of the most promising are selected to move on to Stage IV. At this stage, evaluations are conducted on commercial farms throughout the state of Florida, using clonal field plots with 5 to 45 plants per plot. The final selection consists of genotypes that perform well across years and locations for evergreen or deciduous systems, and are released as cultivars. During the four-stage selection process, genotypes have been visually evaluated using 1 (low) to 5 (high) categorical scores for yield, vigor, healthiness, yield estimated via floral buds, yield estimated via flower numbers, and yield estimated via green berries. Firmness and size have also been visually scored on a scale of 1 (low) to 3 (high). Table 1 shows the average number of samples genotyped and phenotyped across various locations and seasons.

**Table 1.** Mean of the number of observations of eight categorical traits, evaluated over several seasons and breeding stages of the blueberry breeding program, in four Florida regions.

| Trait | Category | North | Citra | Central | South | Seasons | Stages |
| --- | --- | --- | --- | --- | --- | --- | --- |
| Yield | 1 to 5 | 713 | 931 | 67 | 85 | 2014, 2015, 2020, 2021 | II, III, IV |
| Vigor | 1 to 5 | 596 | 119 | 89 | 71 | 2020, 2021, 2022 | III, IV |
| Health | 1 to 5 | 590 | 114 | 85 | 68 | 2020, 2021, 2022 | III, IV |
| Floral Buds | 1 to 5 | 323 | 123 | 89 | 75 | 2021, 2022 | III, IV |
| Flowers | 1 to 5 | 183 | 70 | 89 | 70 | 2021, 2022 | III, IV |
| Green Berries | 1 to 5 | 156 | - | 78 | 60 | 2021, 2022 | III, IV |
| Firmness | 1 to 3 | - | - | 76 | 58 | 2022 | III, IV |
| Size | 1 to 3 | - | - | 76 | 56 | 2022 | III, IV |

For the genomic analyses, we followed the same approach as described in Ferrão, Amadeu, et al. (2021) and Benevenuto et al. (2019). Briefly, genotyping was performed using the “Capture-Seq” approach, and reads were aligned against the largest scaffolds of each of the 12 homoeologous groups of *Vaccinium corymbosum* cv. “Draper” genome assembly (Colle et al., 2019). SNPs were called with FreeBayes v.1.3.2, using 10,000 probe positions as targets (Garrison and Marth, 2012). Loci were filtered, applying the following criteria: (i) minimum mapping quality of 10; (ii) only biallelic locus; (iii) maximum missing data of 50%; (iv) minor allele frequency of 1%; and (v) minimum and maximum mean sequence depth of 3 and 750 across individuals, respectively. A total of 63,552 SNPs were kept after these filtering steps. Sequencing read counts per allele per individual were extracted from the variant call file using vcftools v.0.1.16 (Danecek et al., 2011) and were used as input to estimate the allele dosage, according to the “norm model” in the updog 2.1.0 R package (Gerard et al., 2018).

## Results and Discussion

In this study, we discussed potential scenarios of analysis that underlie a real breeding program, including statistical modeling decisions, forms of data collection, potential errors in evaluation, and traits with different genetic controls. Collectively, we tested three analysis procedures (CONT, CAT-CONT, and CAT), evaluated under three model approaches (Mixed models, Bayesian, and Random Forest), for four number of categories (1 to 2, 1 to 3, 1 to 5, and 1 to 9), with three different noise levels (low, moderate and high) and two hypothetical genetic architectures (qualitative and quantitative). These multiple scenarios create high complexity to present and discuss the results.

To circumvent this, we structured our discussion in the following format. First, we framed our narrative in terms of genomic prediction, and discussed the importance of using different categorical levels to classify continuous traits. Sequentially, we emphasized the impact of using different methods (LMM and GLMM, BLRM and BORM, RFR and RFC) and distributions (continuous and categorical) on predictive accuracy. After discussing the importance of our results for prediction, we focused on inference and considered potential impacts on estimating marker effects, heritability, and genetic gains. Finally, we applied our findings of the simulated populations to a real population of blueberry.

### What is the impact on predicting categorized traits that are continuous by nature?

In breeding programs, phenotypic traits are often recorded using visual scores and categorical traits. We aimed to investigate the impact of categorizing traits, that are continuous by nature. For example, categories ranging from 1 to 5 are commonly used for yield evaluation (Williams et al., 2021), while screening disease progress is often recorded using categories ranging from 1 to 2 (Manichaikul and Broman, 2009) or 1 to 9 (Ferrão, Ferrão, et al., 2019). To answer this question, we tested various categorical levels and different levels of error (Figure 3).

**Figure 3.**
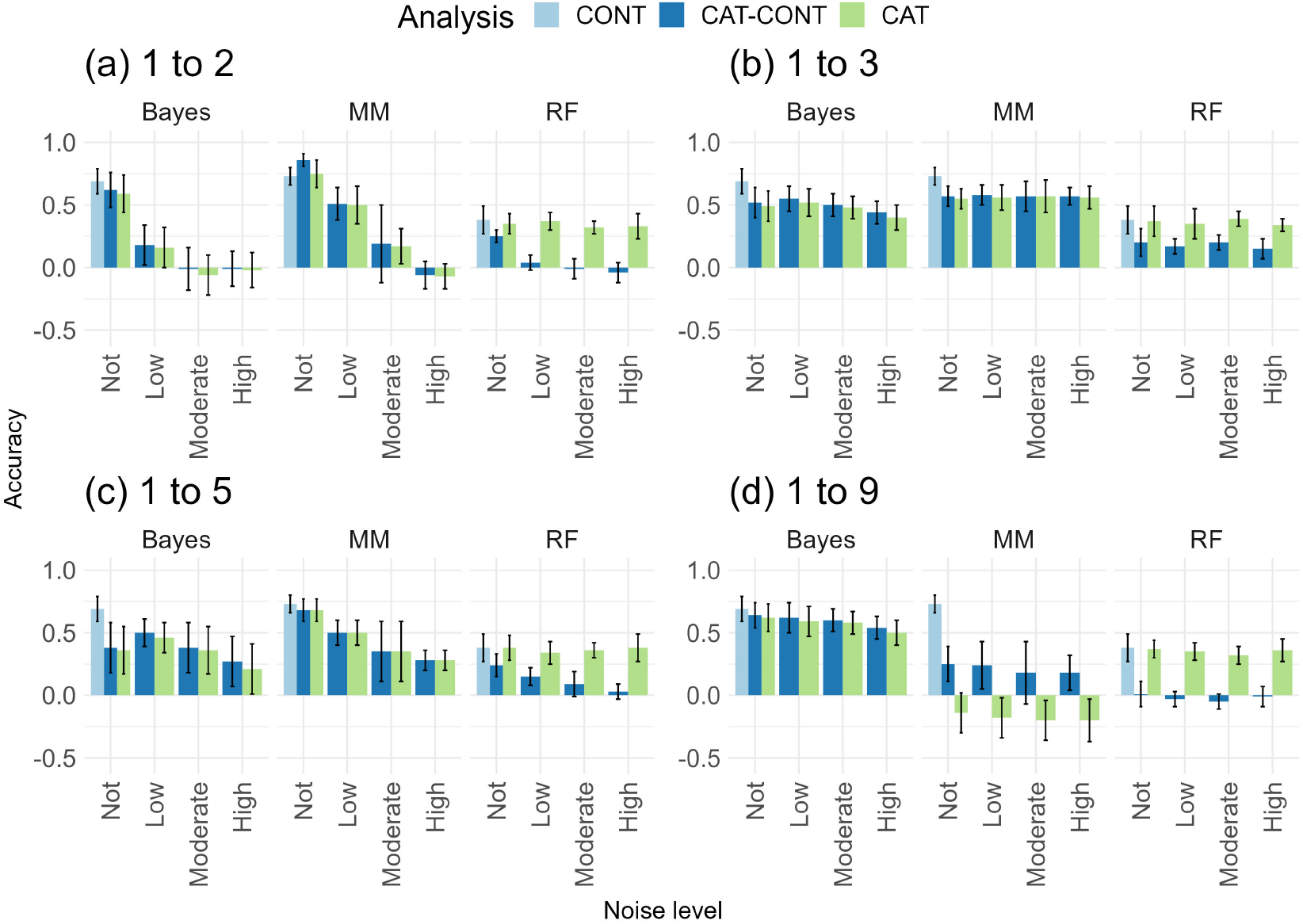
Accuracy of different methods (Bayesian Ordinal and Bayesian Linear Regression Models (Bayes), Generalized and Linear Mixed Models (MM), and Random Forests Regression and Classification (RF)) under cross-validation procedures for categorical traits, with different category levels (1 to 2; 1 to 3; 1 to 5; 1 to 9), under different levels of noise (low - 20% misclassification, moderate - 50% misclassification, and high - 70% misclassification) and no errors, relative to continuous traits. The traits had a quantitative genetic architecture, with 100 QTLs and heritability equal to 0.10.

When compared, scenarios using categories ranging from 1 to 2 were poor simplifications of continuous scenarios (Figure 3a). On the other hand, using more categories (1 to 9) tended to resemble a continuous distribution, although in the presence of larger prediction errors it becomes noisier (Figure 3d). It is reasonable to expect that when more categories are created, we increase misclassification among categories, since scoring extremes led to less subjectivity than assessing intermediate scores. In general, we found that an intermediate number of categories (such as 1 to 3 and 1 to 5) showed a good compromise in terms of predictive accuracy and error, above the other categories classification tested (Figures 3b and 3c).

### LMM and BLRM are robust, but not the best predictive performance for categorical traits

After discussing the relevance of using different numbers of categories to simplify continuous traits, we addressed how breeders should statistically model categorical traits. While GLMM and BORM are more flexible models and have a broader application, their implementation is more computationally demanding, and require a higher level of statistical understanding than standard “normal” methods (LMM and BLRM). Figure 3 shows three scenarios of accuracy evaluation, where breeders need to deal with a combination of strategies, including analyzing categorical data with LMM and BLRM (CAT-CONT), and modeling categorical traits with GLMM and BORM (CAT).

Regarding the simulated genetic architecture, quantitative and qualitative architectures produced very similar results (Figure 3 and Supplemental Figure S1). By focusing primarily on the quantitative results, BLRM and LMM showed the highest accuracies for continuous scale phenotypes (0.69 and 0.73, respectively). When using standard “normal” models to predict a categorical trait (CAT-CONT), we observed reasonable accuracy values (Figure 3). Similarly, Silveira et al. (2019) evaluated rust disease resistance in *Eucalyptus urophylla*, and reported comparable predictive abilities using LMM and GLMM under a Bayesian framework. Ornella et al. (2012) compared parametric and non-parametric models, and reported that LMM had superior prediction performance. Reportedly higher prediction accuracies for the LMM method were published by Heuer et al. (2016), when contrasted with GLMM. In general, our simulated data suggest that the use of BLRM and LMM are robust to ranking genotypes, even when normality assumptions are violated.

While LMM and BLRM may produce consistent results, they may not be the best approach in terms of predictive ability. First, we note that whenever simplification is performed by transforming continuous data into categorical data, there are losses in accuracy. Similarly, Kizilkaya, Fernando, and Garrick (2014) also evaluated continuous and categorical traits, but using Bayes Cπ linear and threshold models, and reported substantially lower accuracies when both traits were compared. A second important aspect relies on the relevance of including noise in the data analyses. When traits are visually assessed, inexperience and subjectivity are both important noise factors to be considered. We observed that RFC was less sensitive to errors, resulting in good predictive accuracy – except for the 1 to 2 category level. Likewise, Biffani et al. (2017) investigated prediction accuracies in mislabeled phenotypic datasets using machine learning methods, and reported similar results. On the relevance of using non-parametric methods, RFC had a better predictive performance than RFR, even in scenarios with a larger number of categories. Therefore, we emphasize that when categorical traits are presented, they should be framed as a classification problem, even if the dataset presents an approximately normal distribution.

Altogether, as an important message for prediction analyses, the use of Gaussian assumptions based on LMM and BLRM for predicting categorical traits, in fact, results in robust prediction values. It provides certain flexibility for breeders and biometricians who are less familiar with GLMM, BORM and machine learning theory. However, in breeding programs where computational runtime is not a problem, marginal gains can be obtained when parametric and non-parametric methods that do not rely on Gaussian assumptions are used (BORM and RFC).

### Genetic parameters are better assessed using Bayesian ordinal regression models

Breeders also rely on the inference of genetic parameters to assist future decisions. For example, most breeding programs have guided their decisions based on the level of genetic control (i.e., heritability), the magnitude of gene action effects, the correlations between traits, and the dynamics of genotype-by-environment interactions. At the molecular level, understanding the genetic architecture of a trait requires estimation of the number, position, and effect size of molecular markers associated with putative QTL.

When we computed the Person’s correlation between marker effects estimated in each scenario (below the diagonal, Figure 4, and Supplemental Figures S2 to S5) for categories ranging from 1 to 5, the correlation values were moderate to high. We also observed a large percentage of agreement between the top 10% of individuals selected across the scenarios (above the diagonal, Figure 4, and Supplemental Figures S2 to S5). Importantly, Bayesian models reported the highest percentage of agreement on selecting the top 10% of individuals, compared to the true breeding value (first row in the matrices, Figure 4, and Supplemental Figures S2 to S5). The concordance between methods when estimating marker effects shed initial light on the robustness of using LMM and BLRM for association analyses (i.e., GWAS) of categorical traits. At the practical level, the large percentage of agreement when ranking genotypes is another important parameter to guarantee genetic gains over generations.

**Figure 4.**
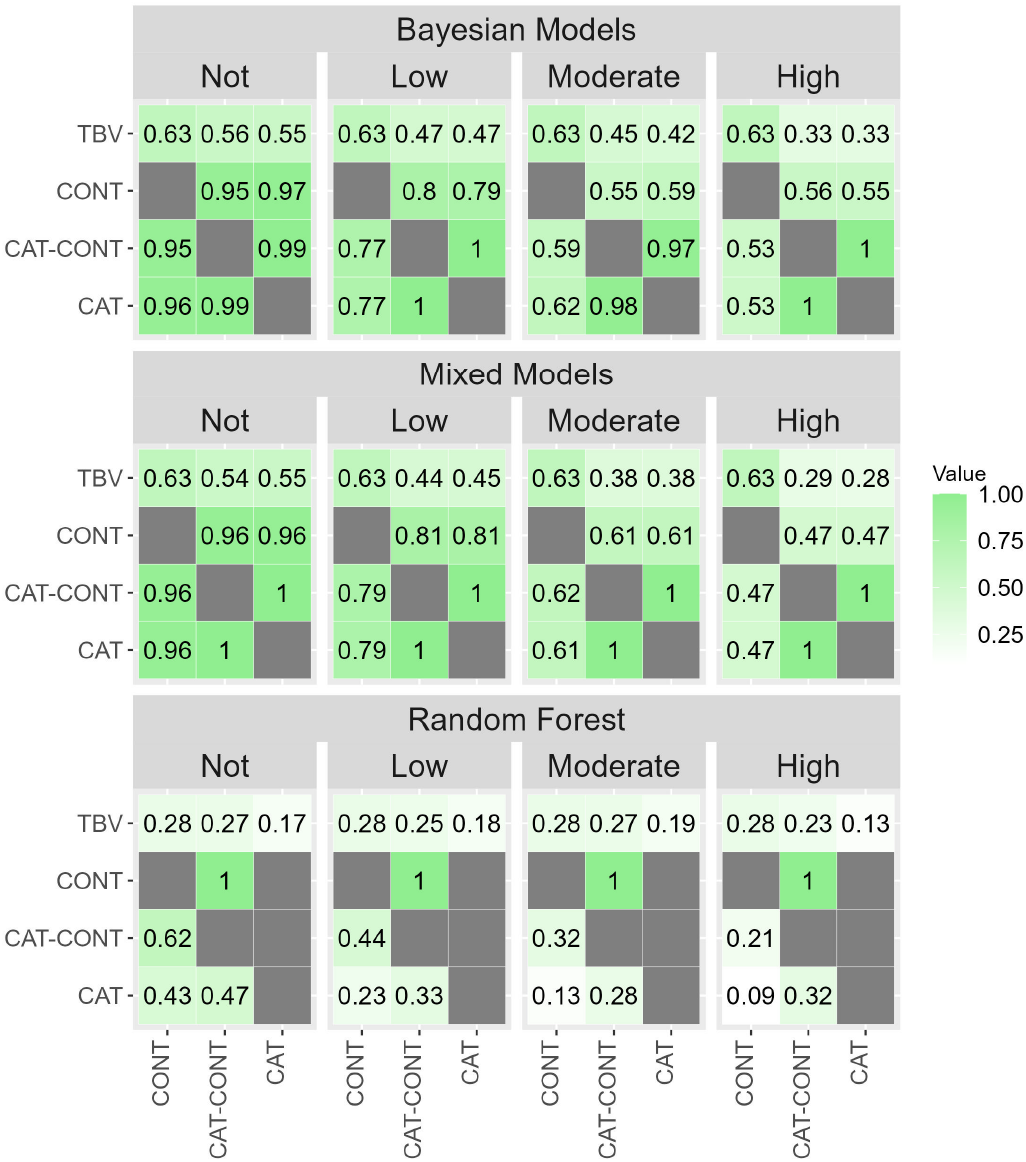
Percentage of agreement between 10% of individuals selected (above the diagonal) for 1 to 5 categorical traits, and correlation between marker effects estimated (below the diagonal) by various analysis approaches (CONT, CAT-CONT and CAT) using Bayesian Ordinal and Bayesian Linear Regression Models, Generalized and Linear Mixed Models, and Random Forests Regression and Classification methods, under different levels of noise (low – 20% of misclassification, moderate – 50% of misclassification, and high – 70% of misclassification) and no errors. The traits had a quantitative genetic architecture, with 100 QTLs and heritability equal to 0.10. The first row in the matrices represents the percentage of agreement between the 10% of individuals selected by the approaches and the true breeding values (TBV).

Another central genetic parameter is heritability. According to Azevedo et al. (2015), recovering heritability values may be more powerful for discriminating between methods, which we used in this study to compare different approaches. We found that on the continuous distribution, Bayesian and mixed models recovered the simulated value of heritability (Figure 5 and Supplemental Figure S6), suggesting that our simulation was appropriate. When contrasting different scenarios, all methods underestimated heritability values. As an important trend, the Bayesian ordinal regression (CAT) presented more robust results in recovering the estimated value. Similar results were also reported by Kizilkaya, Fernando, and Garrick (2014), when using the Bayes Cπ linear model. Tiezzi and Maltecca (2015) evaluated the impact of computing genetic parameters using LMM and GLMM in a Bayesian framework, and found that GLMM captured larger proportions of the genetic variance, resulting in higher heritability values. When comparing parametric and non-parametric methods, the use of RF has some disadvantages, as model covariances between predictors are only possible through recursive expressions, such as the variance of predicted values (Chen and Zhang, 2013). If RF suffers from biased estimates (Supplemental Figures S7 and S8), genetic parameters will also be biased, as we observed in the analyses of the quantitative trait (Figure 5). Finally, we investigated the impact on genetic gain, which relies primarily on heritability values (Supplemental Figures S9 and S10). We found a severe underestimation of genetic gains.

**Figure 5.**
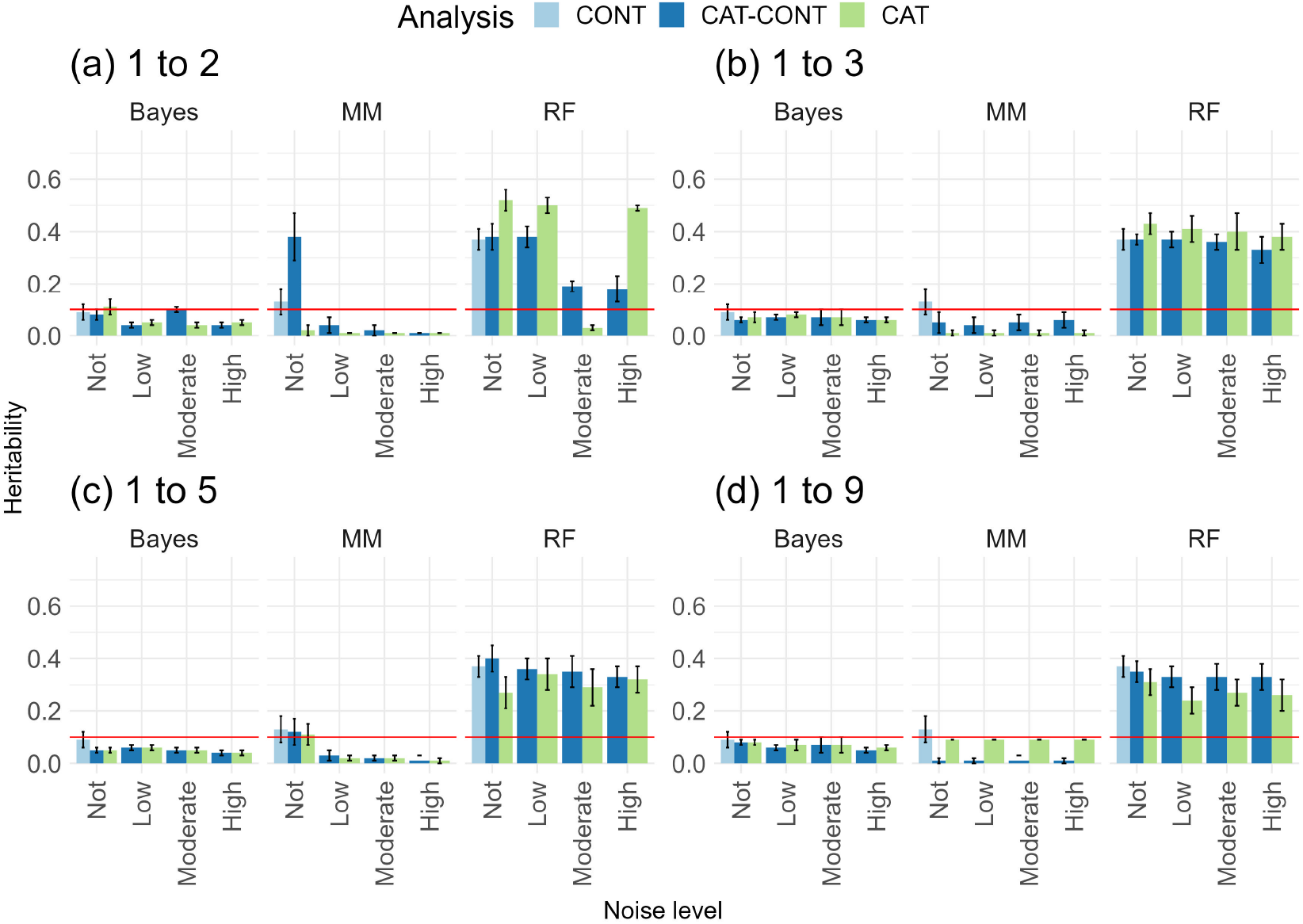
Heritability estimated by different methods (Bayesian Ordinal and Bayesian Linear Regression Models (Bayes), Generalized and Linear Mixed Models (MM), and Random Forests Regression and Classification (RF)) for categorical traits, with different category levels (1 to 2; 1 to 3; 1 to 5; 1 to 9), under different levels of noise (low - 20% misclassification, moderate - 50% misclassification, and high - 70% misclassification) and no errors, relative to continuous traits. The traits had a quantitative genetic architecture, with 100 QTLs and heritability equal to 0.10. The red line represents the simulated heritability value.

### Genomic prediction and inference on the genetic basis of blueberry traits using categorical data

Our next contribution relies on GS implementation using real data. Along the years, blueberry breeders often face the dilemma of phenotyping a large breeding population using visual scores, or focusing on a restricted number of samples and collect accurate continues phenotypes. In this context, yield is a prime example. While blueberry bushes need to be harvested multiple times over the season, the phenotyping process is labor intensive, costly, and low throughput. To circumvent this, the UF blueberry breeding program has the following strategy: berries from mature plants on advantaged breeding stages are manually harvested and weighed, while plants from earlier stages are visually scored based on general yield, number of flowers, flowers buds and green fruits. While continuous yield data are formally used to train predictive models for genomic selection, visual scores are used as auxiliary traits, and are used by breeders to make decisions after ranking the genotypes, using main fruit quality traits and yield.

The prediction results detected in the simulated data encouraged us to explore genomic prediction in blueberry from three different angles. First, we computed predictive ability and genetic parameters for multiple categorical traits, and observed that the Bayesian model showed the highest values (Figure 6 and Supplemental Table S1). The Bayesian approach has also demonstrated more reasonable values for heritability, in previous research. For example, using pedigree and continuous data, Cellon et al. (2018) reported ∼ 0.4 and ∼ 0.3 narrow-sense heritability values for firmness and size in blueberry, respectively. Similar values were also reported by de Bem Oliveira et al. (2020), but using molecular information.

**Figure 6.**
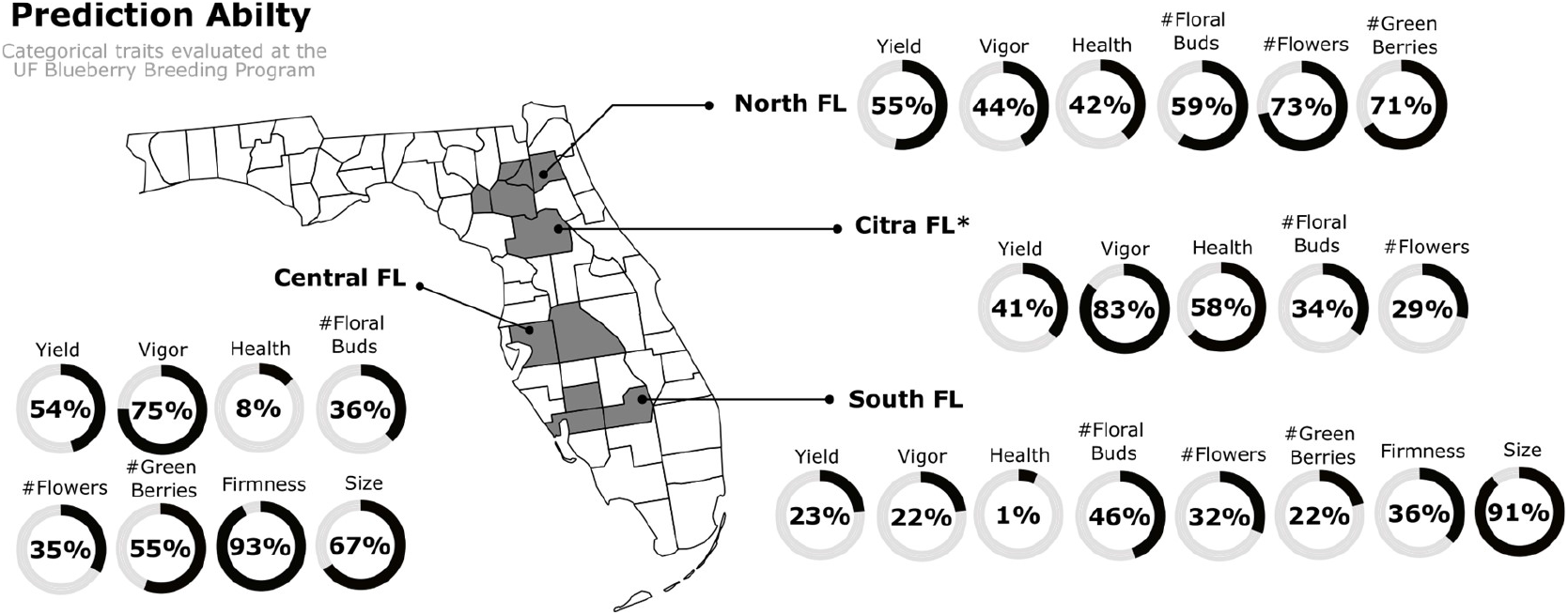
Genomic prediction was performed using cross-validation procedures. The mean predictive ability using categorical phenotypes collected in several seasons (2014, 2015, 2020 to 2022) and breeding stages (II, III, and IV) over four macro-regions in Florida State. All predictive abilities were expressed as percentage values.

In the sequence, we explored the underlying question of the advantage of increasing the population training size for genomic prediction, when continuous and categorical data are used. For integrating categorical and continuous data, we identified a group of 179 samples that were both phenotyped for yield (kg), in a continuous and categorical form. The estimated heritability using continuous data was 0.28, and a similar value was computed using categorical data (Supplemental Table S1). However, substantial differences were observed for prediction, when yield was considered as continuous and categorical data. Yield prediction for the 179 genotypes using continuous distribution and cross-validation scheme was 0.17. To test the relevance of leveraging predictive ability at the cost of including more visually scored individuals, we trained our model using a new set of 2,323 genotypes, all visually scored using categories ranging from 1 to 5. This new set of genotypes was used to predict the 179 genotypes collected at the continuous distribution. Remarkably, predictive ability increased to 0.35, representing more than a 100% gain over the continuous metric. This fact leads to us raising our last scientific question: what is the best alternative to combine continuous and categorical data in a single framework for genomic prediction?

To answer this question, we used the same stochastic process to simulate traits, but under four different genetic architectures (Figure 7). By mimicking our breeding program, we simulated 200 phenotypes in a continuous distribution. Additionally, new genotypes (ranging from 0 to 4,800 with an increment number of 200 individuals) were categorized with different error levels. Herein, our benchmark is the predictive ability computed, using the 5,000 continuous phenotypes. Systematically, we included phenotypes collected at a categorical distribution, and checked the predictive ability (Figure 7). As might be expected, genetic architecture and noise levels are main drivers of predictive ability, with simple genetic architecture and low levels of errors leading to higher predictive ability. When no errors are simulated, increasing the population size by using more categorical traits will always increase the predictive ability of continuous traits. This is an ideal, but unrealistic scenario. A more realistic approach is a breeding program operating with low noise level. Herein, including more categorial phenotypes only adds to predictive ability when error levels are low, a fact that sheds light on the importance of properly training for data collection.

**Figure 7.**
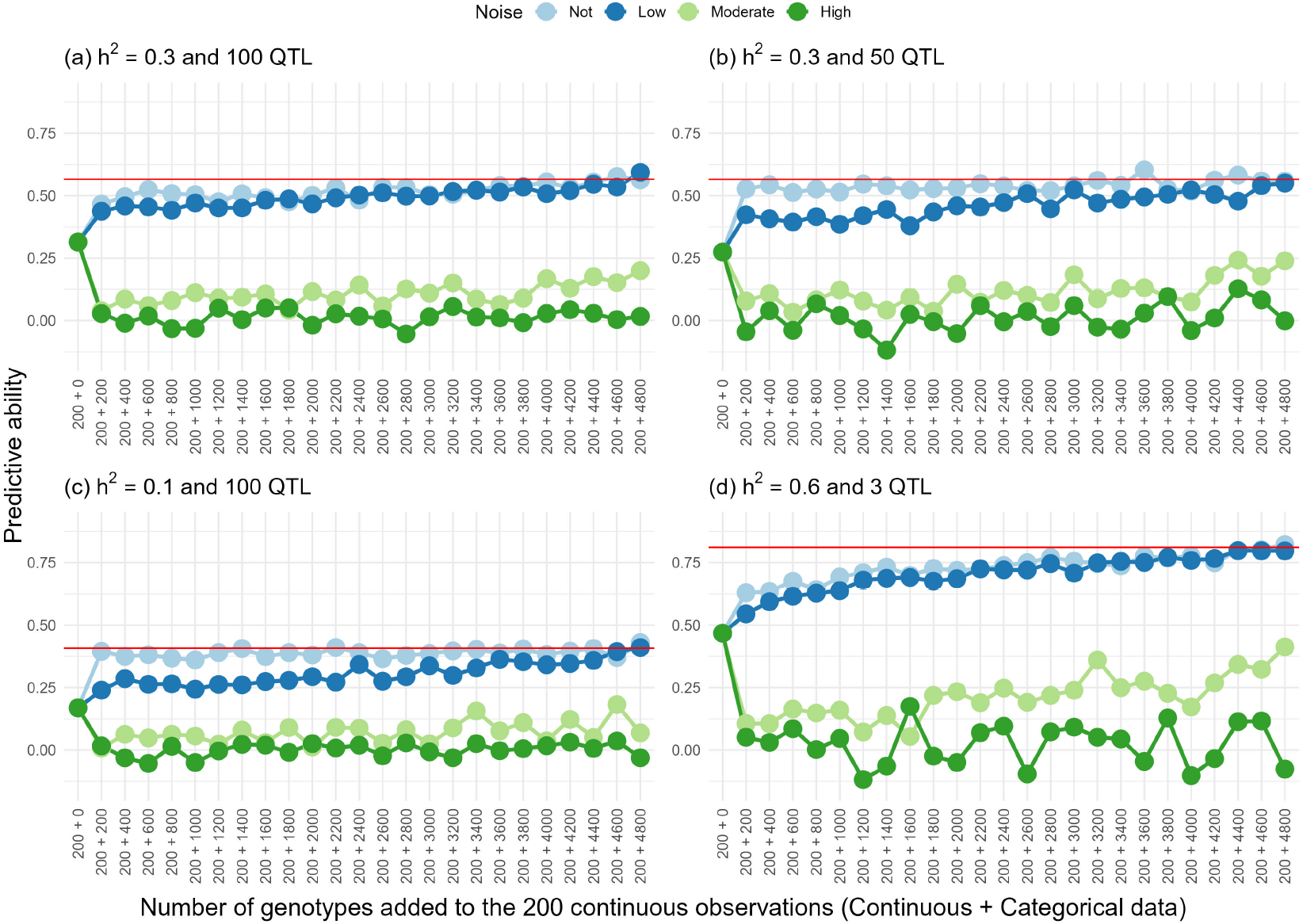
Genomic prediction was performed using cross-validation procedures for categorical traits, considering categories ranging from 1 to 5, different levels of noise (low – 20% of misclassification, moderate – 50% of misclassification, and high – 70% of misclassification), and no errors. The genetic architectures of the traits were: (a) h2=0.30 and 100 QTLs, (b) h2=0.30 and 50 QTLs, (c) h2=0.10 and 100 QTLs, and (d) h2=0.60 and 3 QTLs. The predictive abilities were calculated using different numbers of genotypes (ranging from 0 to 4,800 in increments of 200) recorded in the categorical trait, in addition to the 200 genotypes recorded in the continuous distribution. The horizontal red line represents the predictive ability of 5,000 genotypes recorded in the continuous distribution in each scenario.

## Conclusion

Altogether, the real and simulated data used in this investigation allow us to provide a blueprint for how visual scores could be used by plant breeders. Regarding our main research questions, we can first conclude that categorical traits can be effectively used for prediction and inference on traits with different genetic architecture, with gains and precision directly related to the amount of noise and subjectivity included in the analyses. Secondly, the use of traditional statistical approaches showed robustness, but not the best predictive results over different error levels and genetic architectures. Thus, when time and computational resources are not a barrier, the use of Bayesian ordinal regression models are preferable. Next, we reported large predictive abilities for a group of categorical traits collected in blueberry, which opens important venues to include such traits in a molecular breeding pipeline. Finally, we integrated continuous and categorical data and simulated scenarios of genomic prediction for traits with different genetic architectures. Simply stated, we suggested that by including 600-1,000 categorical data phenotypes with low error, we can verify the improvement of stable predictive performance. At this point, breeders are encouraged to reflect on the importance of allocating resources to training their team and the costs related to phenotyping. In the case of blueberry, it is noteworthy that collecting yield and other fruit set traits over the seasons is costly and time-consuming. Investing in better training associated with visual scores is a more feasible alternative when phenotyping is designed for larger populations – at least until the use of computer vision methods for high-throughput phenotyping is fully adopted.

## Acknowledgment

This work was supported by the UF royalty fund generated by the licensing of blueberry cultivars. This preprint was created using the LaPreprint template (https://github.com/roaldarbol/lapreprint) by Mikkel Roald-Arbøl.

## Author contributions

PM, LF and CA conceived and supervised the study. JB coordinated the collection and genotyping of the samples. LF, CA, MDV, MN and ACCN analyzed and interpreted the genomic selection results. LF and CA wrote the paper and included the revision from all authors. All authors read, reviewed and approved the final version of the manuscript for publication.

## Supplementary

The following are the Supplementary Table and Figures to this article:

**Table S1.**
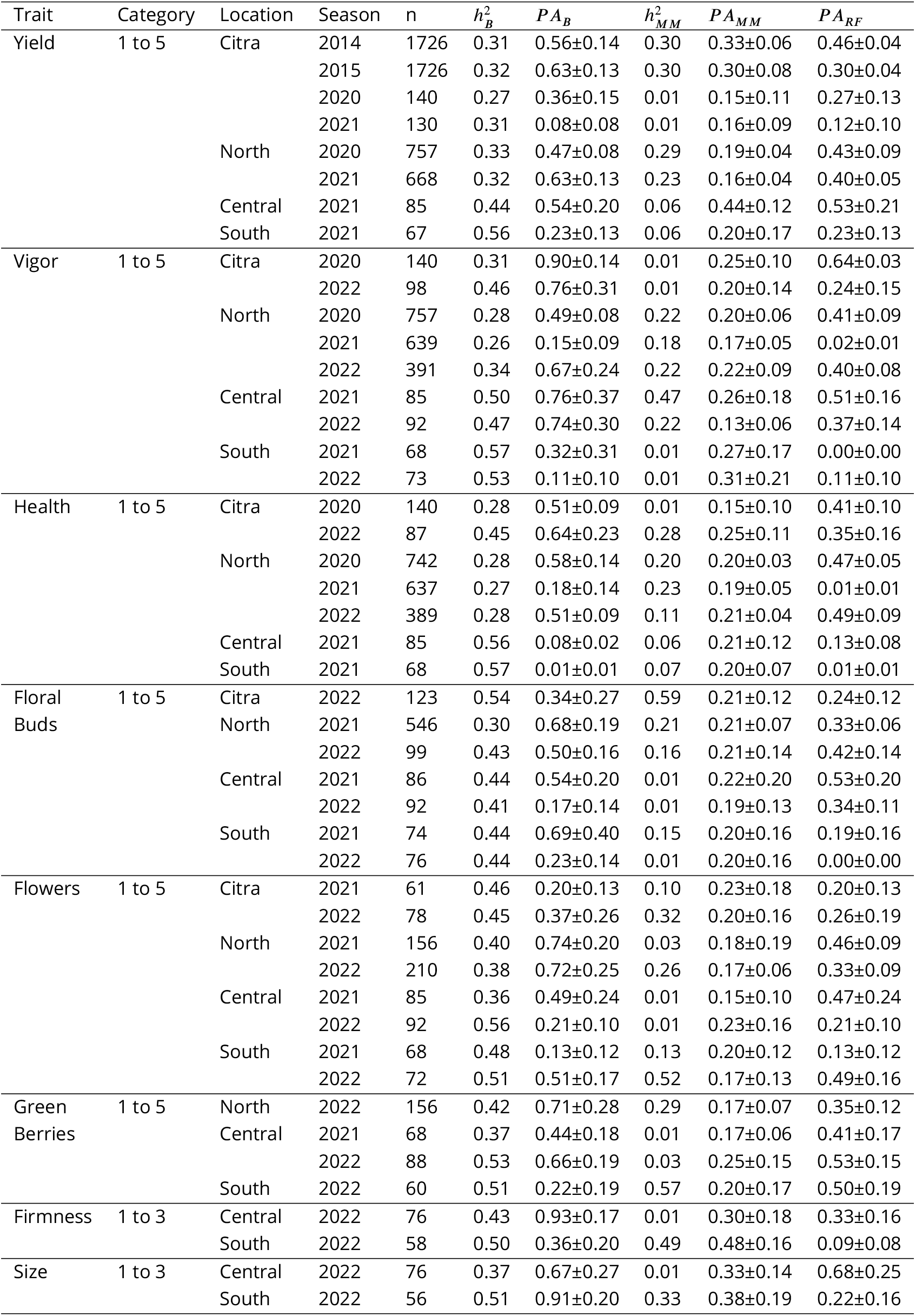
Number of observations (*n*), predictive abilities (*P A*_*i*_) mean and standard deviation and heritability estimates 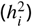, considering the ith method (B: Bayesian, MM: Mixed model, and RF: Random Forest) of eight categorical traits, evaluated in several seasons and stages of the blueberry breeding program, in four Florida regions.

**Figure S1.**
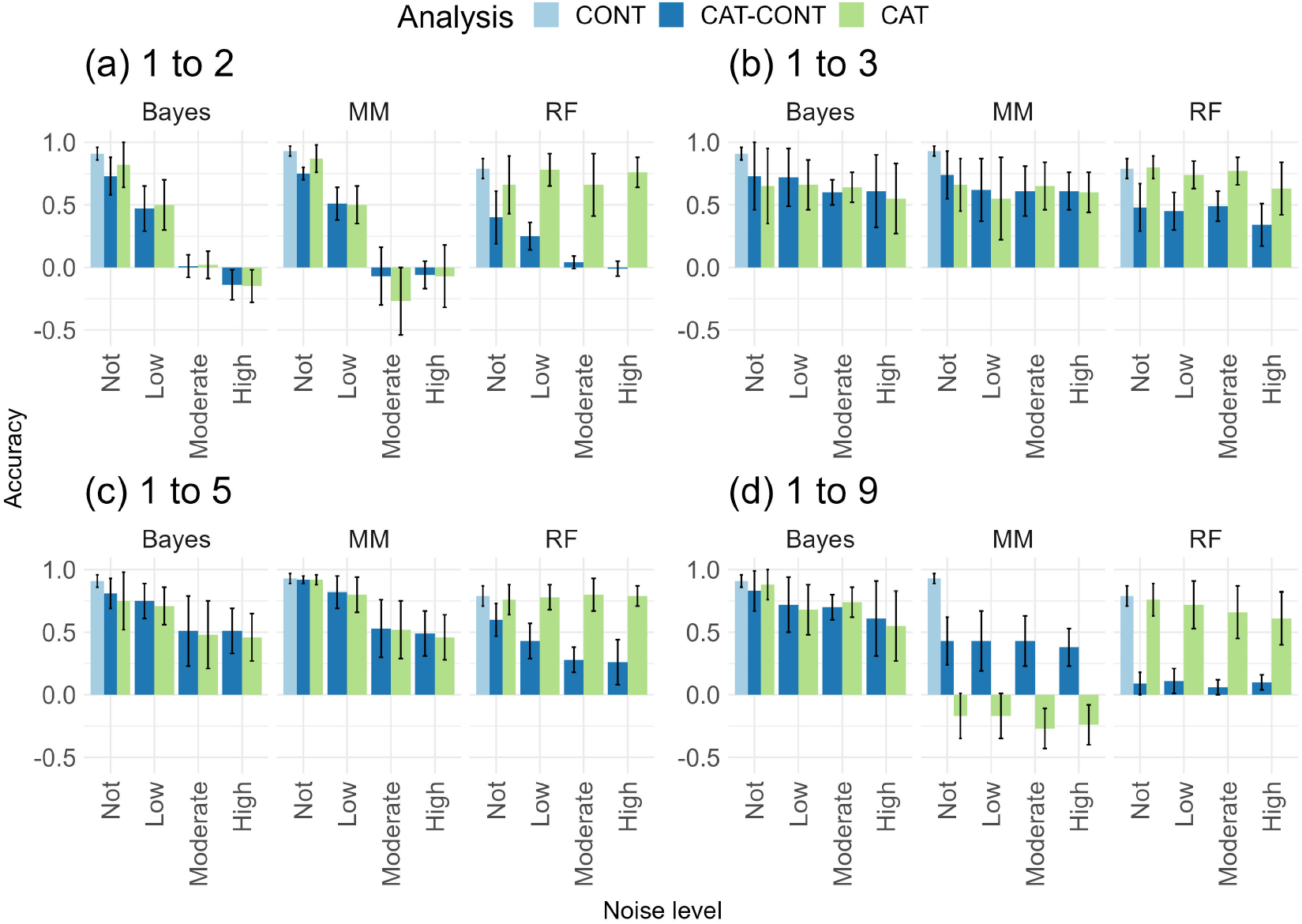
Accuracy of different methods (Bayesian Ordinal and Bayesian Linear Regression Models (Bayes), Generalized and Linear Mixed Models (MM), and Random Forests Regression and Classification (RF)) under cross-validation procedures for categorical traits, with different category levels (1 to 2; 1 to 3; 1 to 5; 1 to 9), under different levels of noise (low - 20% misclassification, moderate - 50% misclassification, and high - 70% misclassification) and no errors, relative to a continuous traits. The traits had a qualitative genetic architecture with 3 QTLs and heritability equal to 0.60.

**Figure S2.**
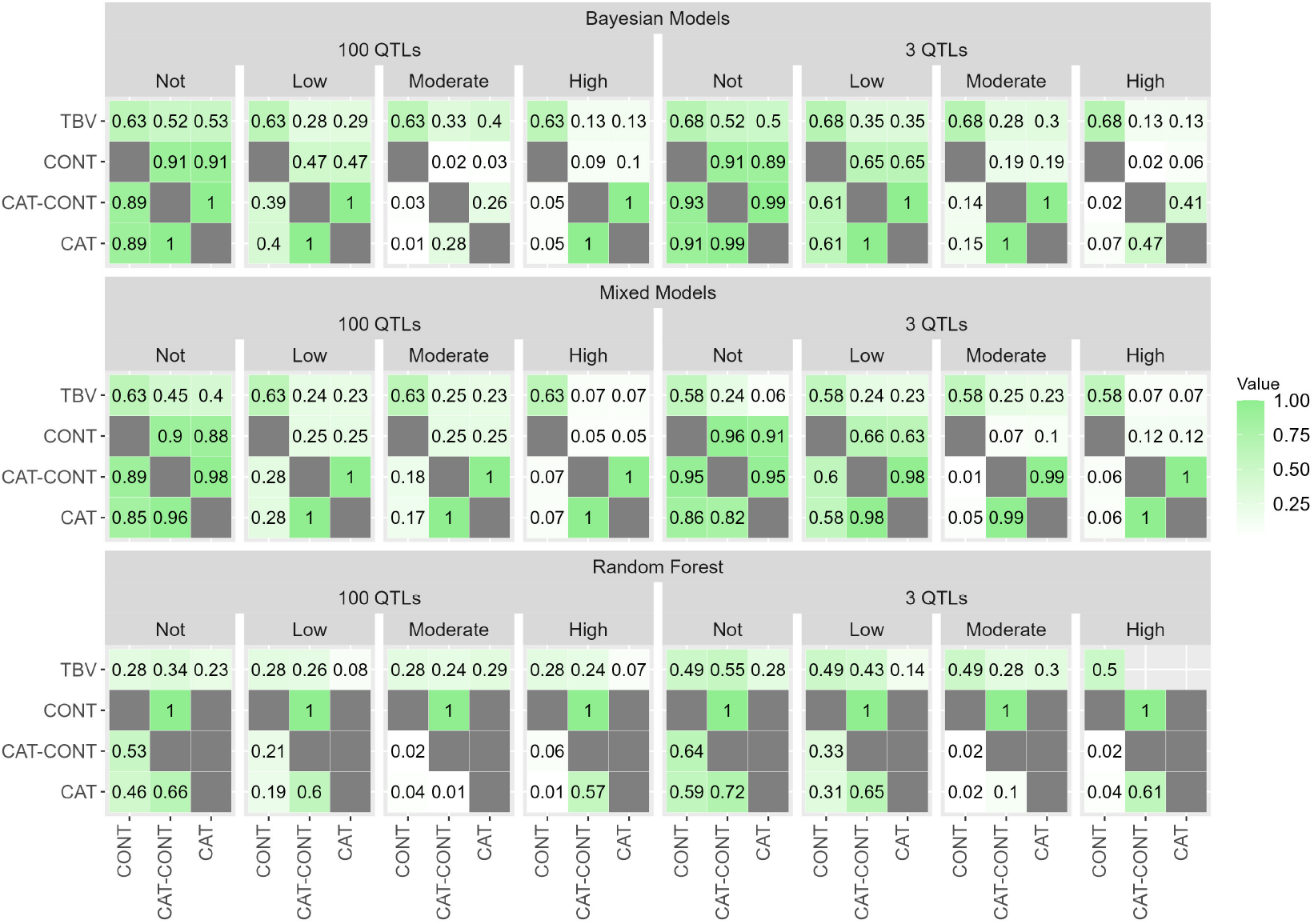
Percentage of agreement between 10% of individuals selected (above the diagonal) for categorical traits with 1 to 2 category levels, and correlation between the marker effects estimated (below the diagonal) by different analysis approaches (CONT, CAT-CONT, and CAT) using Bayesian Ordinal and Bayesian Linear Regression Models, Generalized and Linear Mixed Models, and Random Forests Regression and Classification methods, under different levels of noise (low – 20% of misclassification, moderate – 50% of misclassification, and high – 70% of misclassification) and no errors. The traits had quantitative and qualitative genetic architectures with 100 QTLs and heritability equal to 0.10 and 3 QTLs and heritability equal to 0.60, respectively. The first row in the matrices represents the percentage of agreement between 10% of individuals selected by the approaches and the true breeding values (TBV).

**Figure S3.**
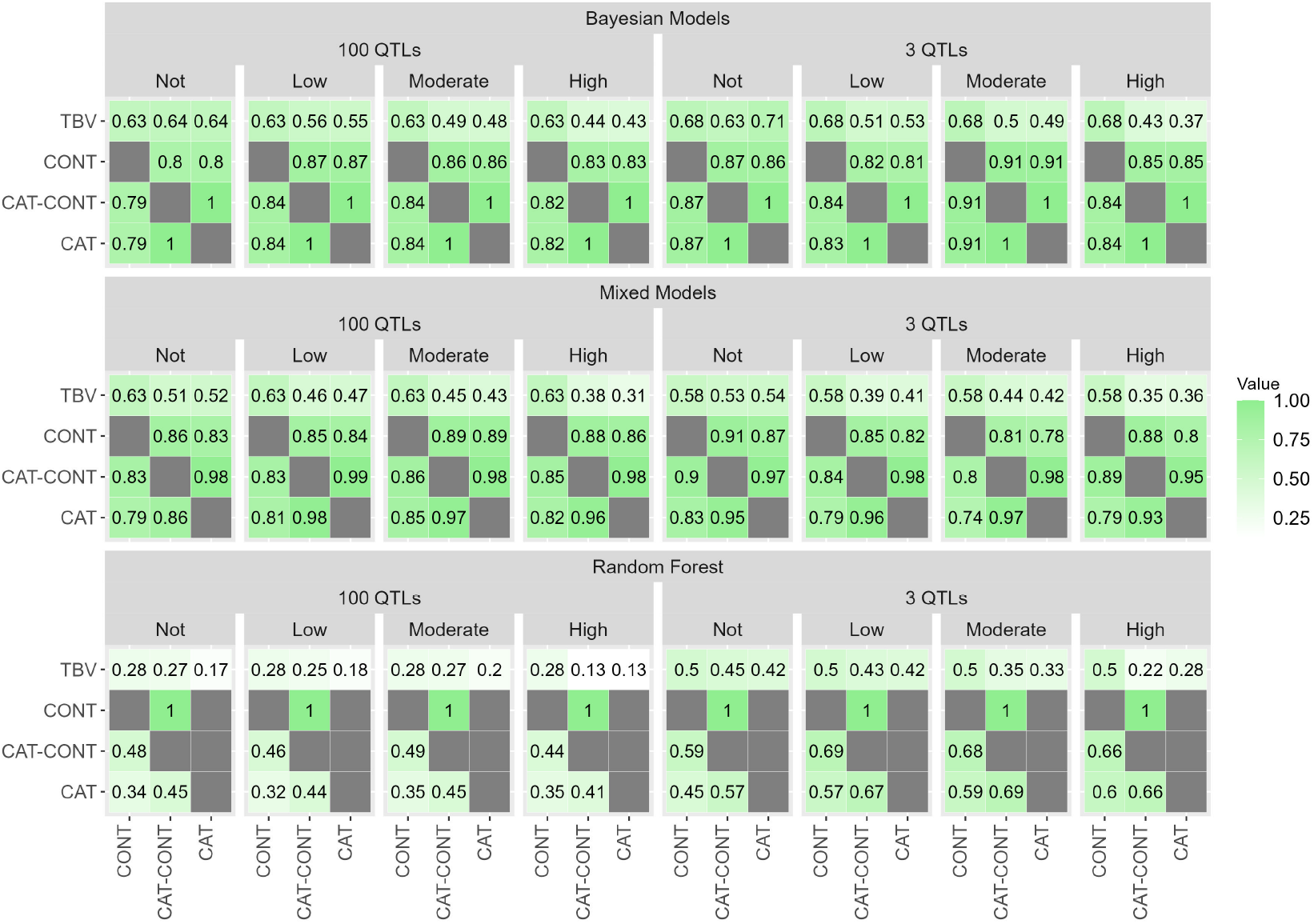
Percentage of agreement between 10% of individuals selected (above the diagonal) for categorical traits with 1 to 3 category levels, and correlation between the marker effects estimated (below the diagonal) by different analysis approaches (CONT, CAT-CONT, and CAT) using Bayesian Ordinal and Bayesian Linear Regression Models, Generalized and Linear Mixed Models, and Random Forests Regression and Classification methods, under different levels of noise (low – 20% of misclassification, moderate – 50% of misclassification, and high – 70% of misclassification) and no errors. The traits had quantitative and qualitative genetic architectures, with 100 QTLs and heritability equal to 0.10, and 3 QTLs and heritability equal to 0.60, respectively. The first row in the matrices represents the percentage of agreement between 10% of individuals selected by the approaches and the true breeding values (TBV).

**Figure S4.**
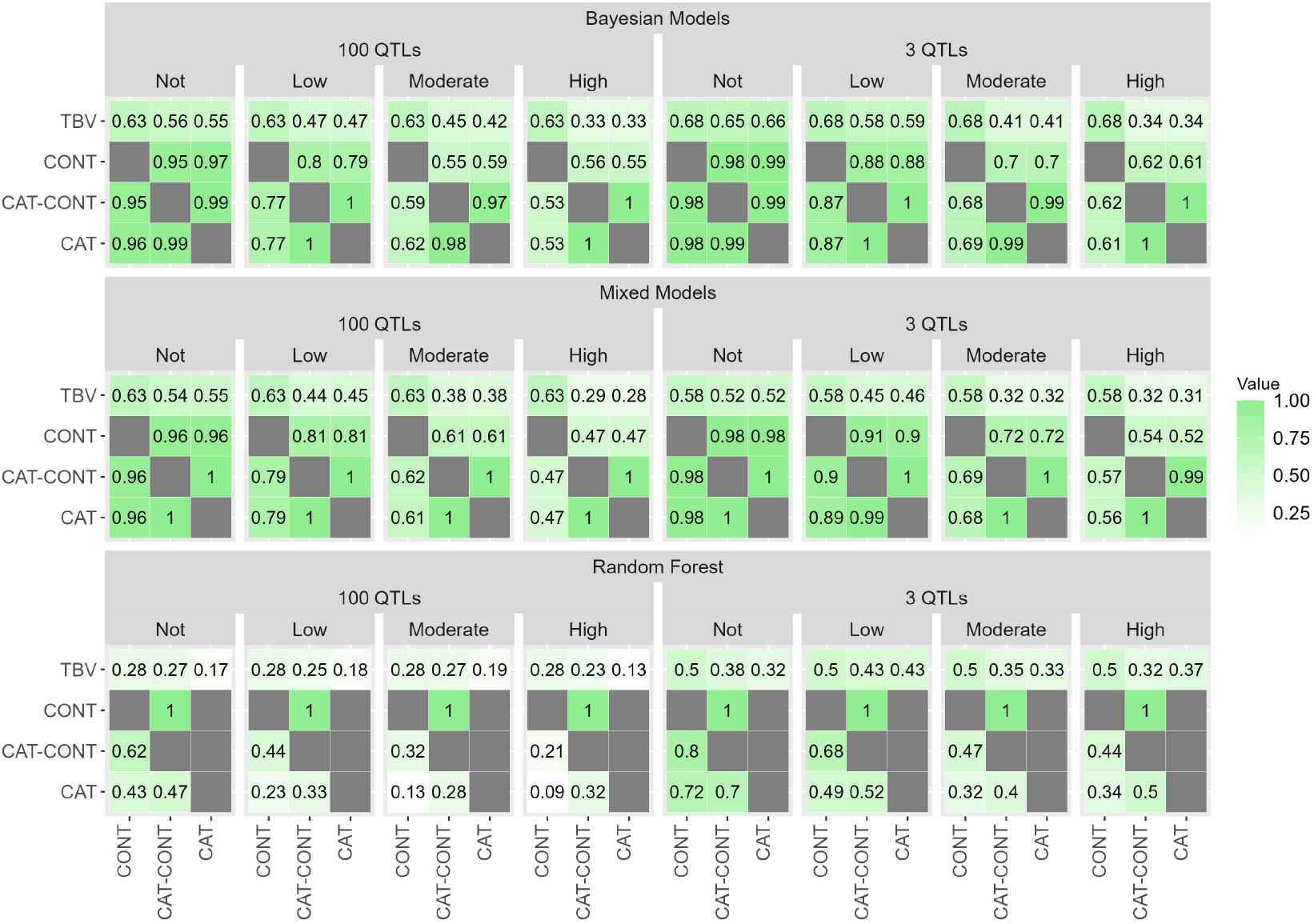
Percentage of agreement between 10% of individuals selected (above the diagonal) for categorical traits with 1 to 5 category levels, and correlation between the marker effects estimated (below the diagonal) by different analysis approaches (CONT, CAT-CONT, and CAT) using Bayesian Ordinal and Bayesian Linear Regression Models, Generalized and Linear Mixed Models, and Random Forests Regression and Classification methods, under different levels of noise (low – 20% of misclassification, moderate – 50% of misclassification, and high – 70% of misclassification) and no errors. The traits had quantitative and qualitative genetic architectures, with 100 QTLs and heritability equal to 0.10, and 3 QTLs and heritability equal to 0.60, respectively. The first row in the matrices represents the percentage of agreement between 10% of individuals selected by the approaches and the true breeding values (TBV).

**Figure S5.**
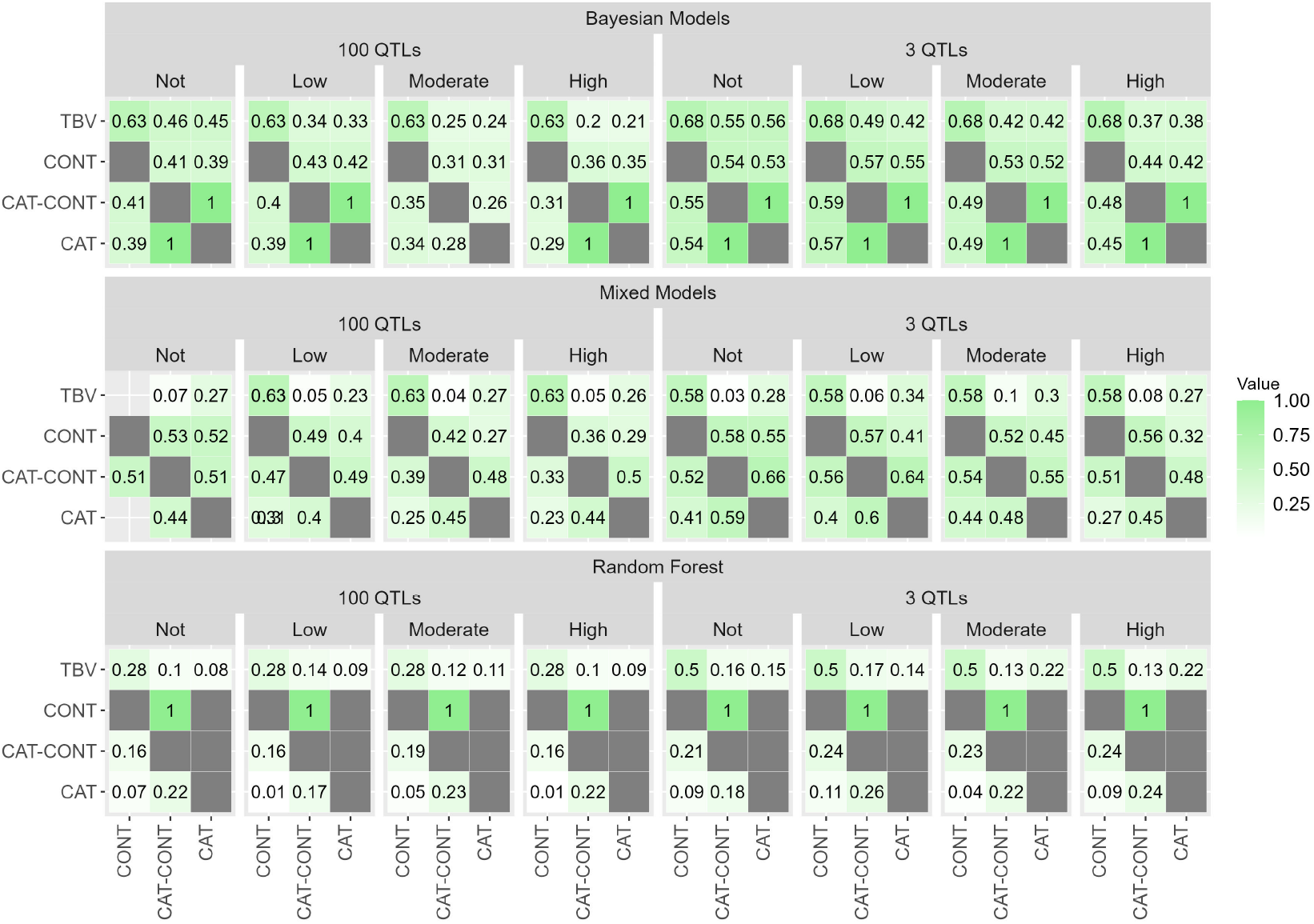
Percentage of agreement between 10% of individuals selected (above the diagonal) for categorical traits with 1 to 9 category levels, and correlation between the marker effects estimated (below the diagonal) by different analysis approaches (CONT, CAT-CONT, and CAT) using Bayesian Ordinal and Bayesian Linear Regression Models, Generalized and Linear Mixed Models, and Random Forests Regression and Classification methods, under different levels of noise (low – 20% of misclassification, moderate – 50% of misclassification, and high – 70% of misclassification) and no errors. The traits had quantitative and qualitative genetic architectures, with 100 QTLs and heritability equal to 0.10, and 3 QTLs and heritability equal to 0.60, respectively. The first row in the matrices represents the percentage of agreement between 10% of individuals selected by the approaches and the true breeding values (TBV).

**Figure S6.**
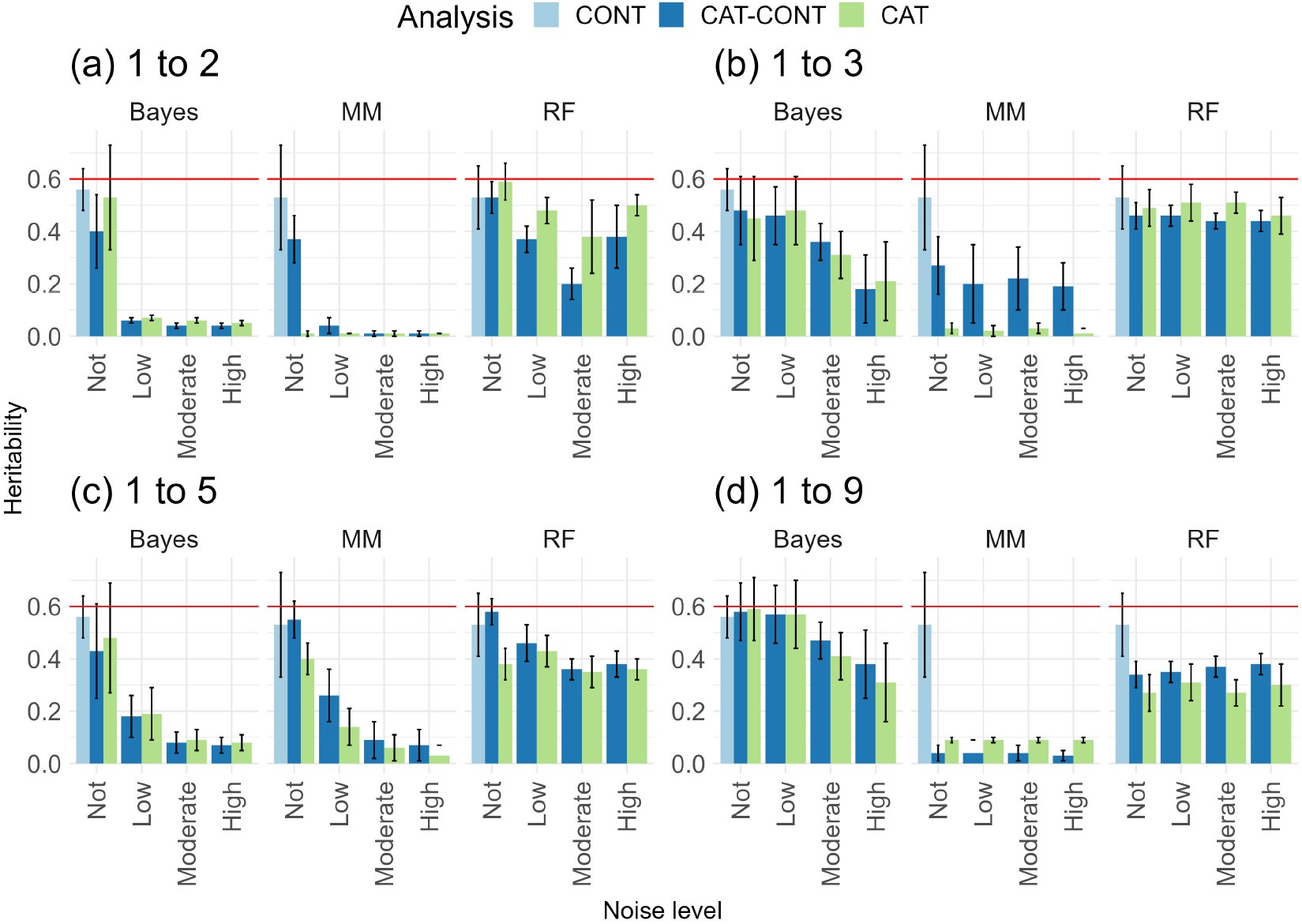
Heritability estimated by different methods (Bayesian Ordinal and Bayesian Linear Regression Models (Bayes), Generalized and Linear Mixed Models (MM), and Random Forests Regression and Classification (RF)) for categorical traits, with different category levels (1 to 2; 1 to 3; 1 to 5; 1 to 9), under different levels of noise (low - 20% misclassification, moderate - 50% misclassification, and high - 70% misclassification) and no errors, relative to a continuous traits. The traits had a qualitative genetic architecture, with 3 QTLs and heritability equal to 0.60. The red line represents the simulated heritability value.

**Figure S7.**
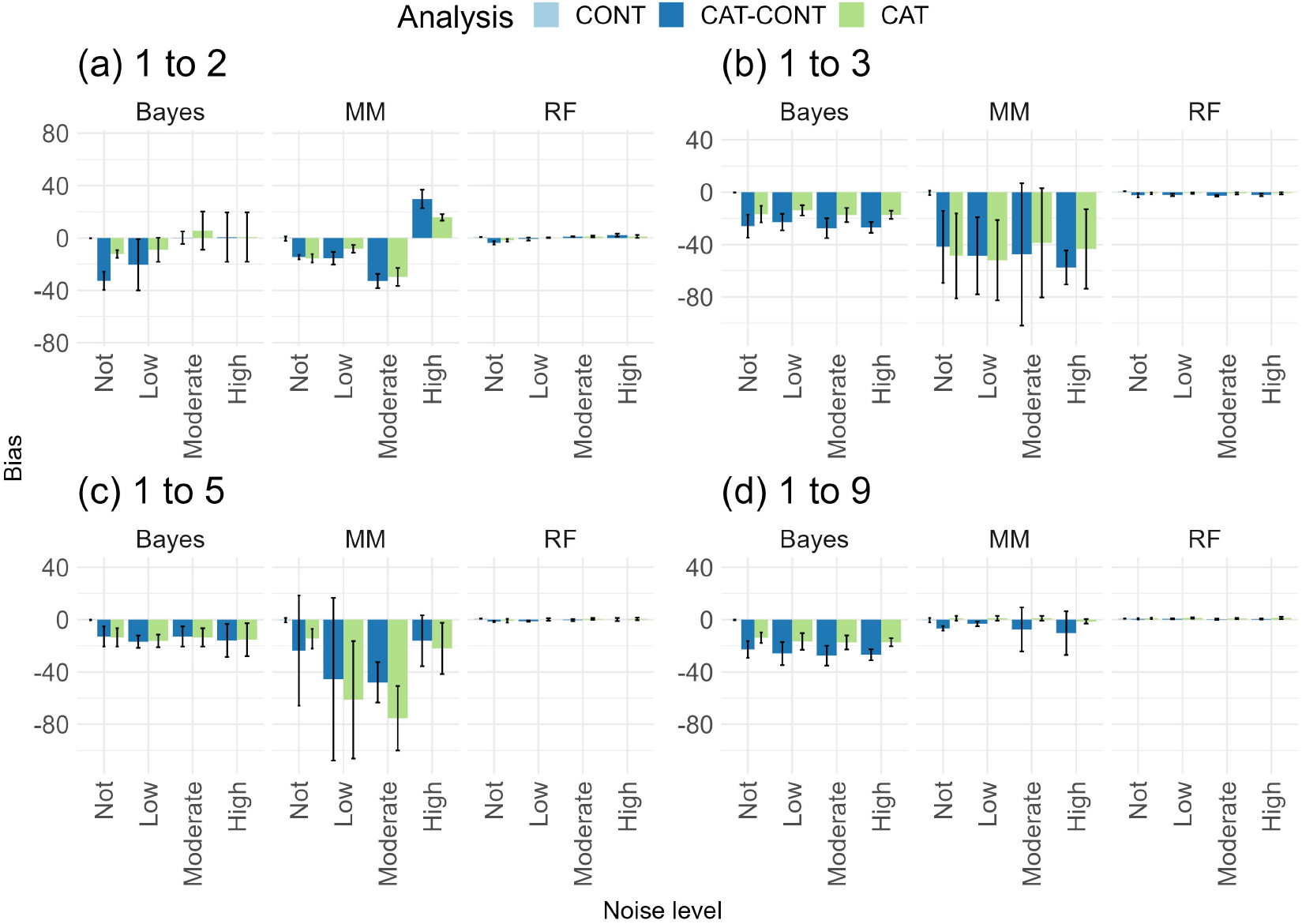
Bias of genomic estimated breeding values from different methods (Bayesian Ordinal and Bayesian Linear Regression Models (Bayes), Generalized and Linear Mixed Models (MM), and Random Forests Regression and Classification (RF)) for categorical traits, with different category levels (1 to 2; 1 to 3; 1 to 5; 1 to 9), under different levels of noise (low – 20% misclassification, moderate – 50% misclassification, and high – 70% misclassification) and no errors, relative to a continuous traits. The traits had a quantitative genetic architecture, with 100 QTLs and heritability equal to 0.10.

**Figure S8.**
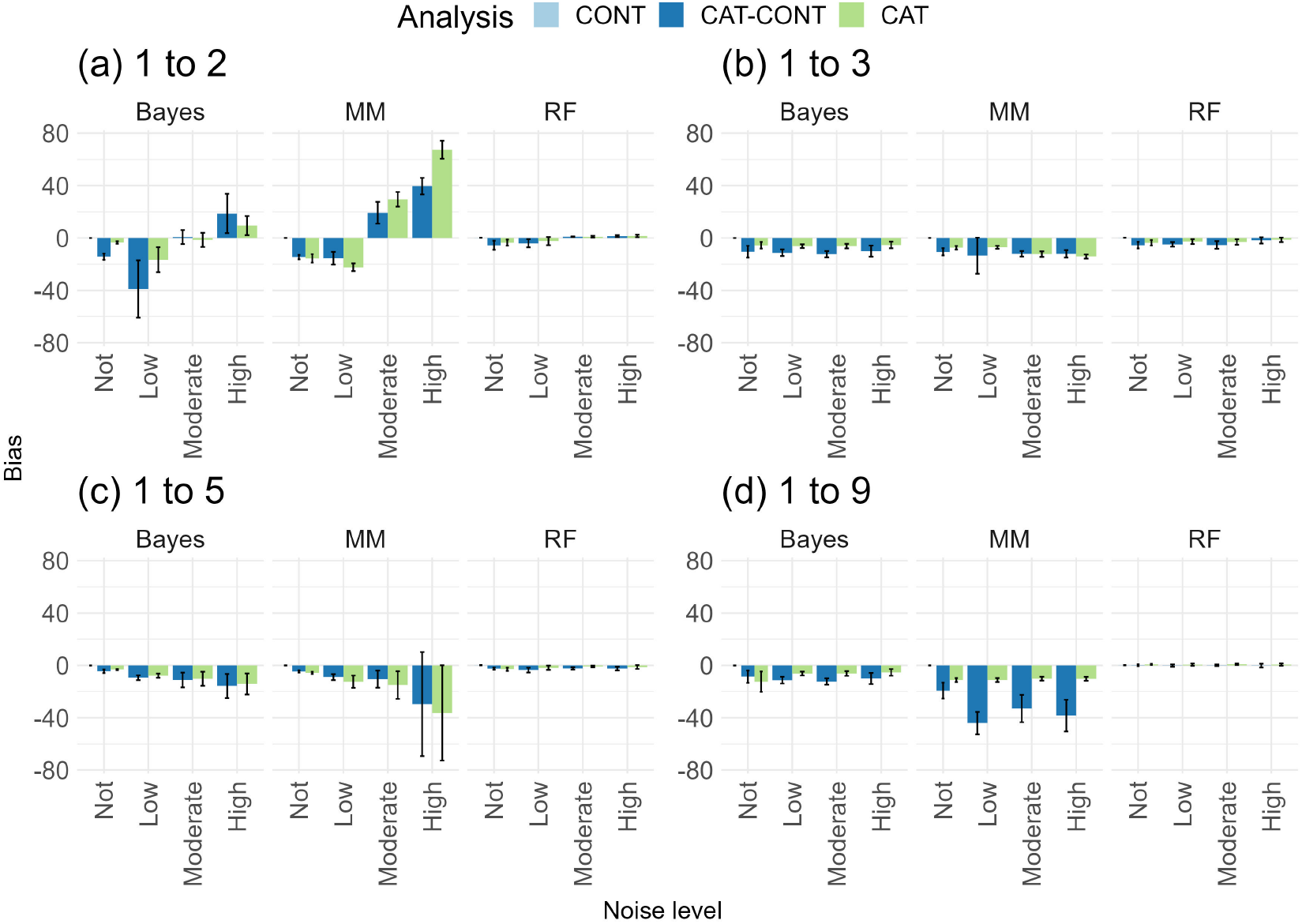
Bias of genomic estimated breeding values from different methods (Bayesian Ordinal and Bayesian Linear Regression Models (Bayes), Generalized and Linear Mixed Models (MM), and Random Forests Regression and Classification (RF)) for categorical traits, with different category levels (1 to 2; 1 to 3; 1 to 5; 1 to 9), under different levels of noise (low - 20% misclassification, moderate - 50% misclassification, and high - 70% misclassification) and no errors, relative to a continuous traits. The traits had a qualitative genetic architecture, with 3 QTLs and heritability equal to 0.60.

**Figure S9.**
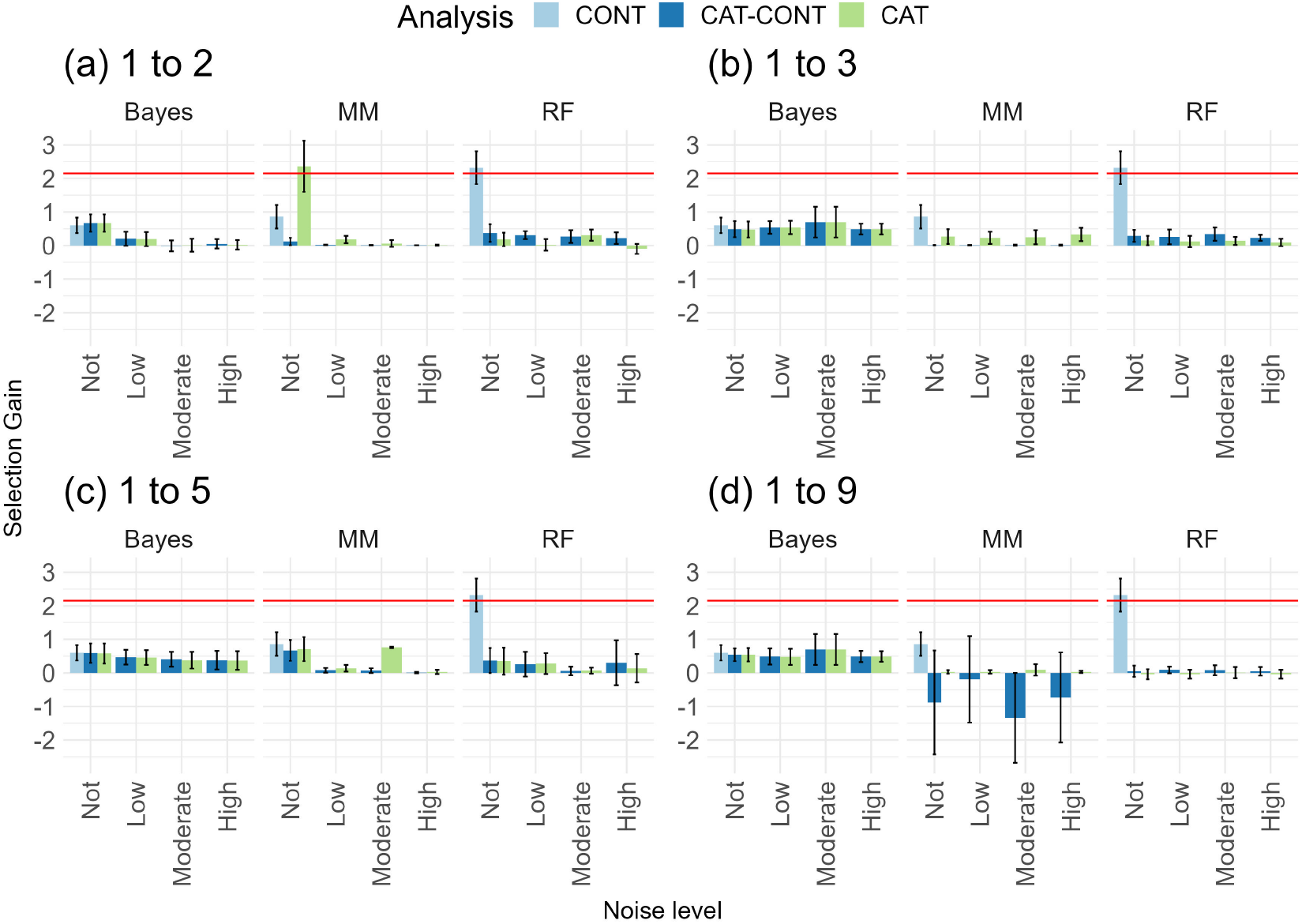
Selection gain estimated by different methods (Bayesian Ordinal and Bayesian Linear Regression Models (Bayes), Generalized and Linear Mixed Models (MM), and Random Forests Regression and Classification (RF)) for categorical traits, with different category levels (1 to 2; 1 to 3; 1 to 5; 1 to 9), under different levels of noise (low - 20% misclassification, moderate - 50% misclassification, and high - 70% misclassification) and no errors, relative to a continuous traits. The traits had a quantitative genetic architecture, with 100 QTLs and heritability equal to 0.10. The red line represents the simulated selection gain value.

**Figure S10.**
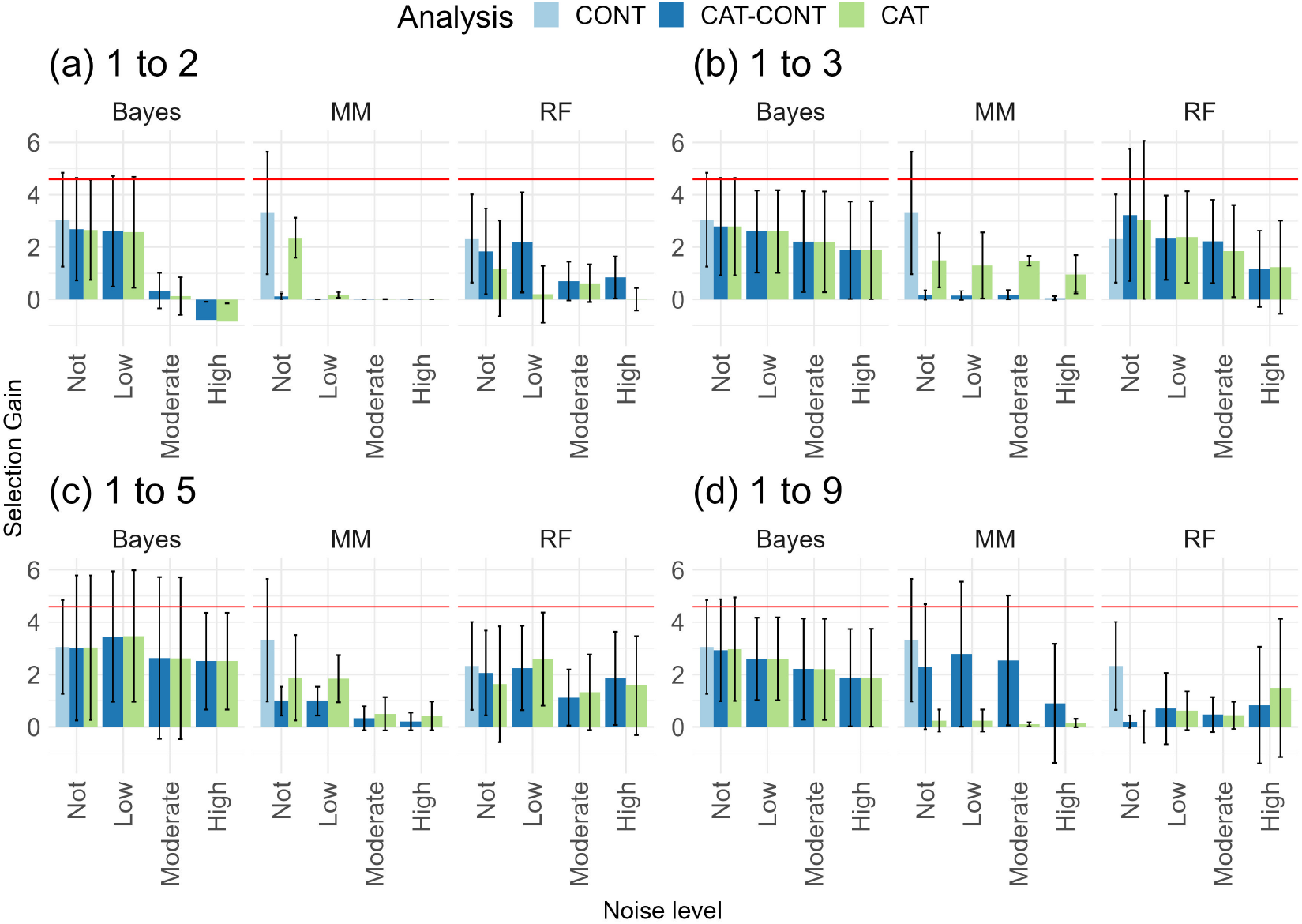
Selection gain estimated by different methods (Bayesian Ordinal and Bayesian Linear Regression Models (Bayes), Generalized and Linear Mixed Models (MM), and Random Forests Regression and Classification (RF)) for categorical traits, with different category levels (1 to 2; 1 to 3; 1 to 5; 1 to 9), under different levels of noise (low - 20% misclassification, moderate - 50% misclassification, and high - 70% misclassification) and no errors, relative to a continuous traits. The traits had a qualitative genetic architecture, with 3 QTLs and heritability equal to 0.60. The red line represents the simulated selection gain value.

